# Safety Assessment and Evaluation of Novel Thiourea Derivatives as Antivirals

**DOI:** 10.1101/2023.06.25.546447

**Authors:** Jitendra Kumar, Purnima Tyagi, Akhilesh K Saini, Deepti Sharma, Vijay Kumar

## Abstract

DSA-00 and its two derivatives (DSA-02 & DSA-09) are new thiourea derivatives that exhibit strong antiviral activity against hepatitis B virus comparable to Entecavir. Here these compounds were evaluated for *in-vitro* cytotoxicity and genotoxicity, and *in vivo* acute toxicity for their potential therapeutic use. The cytotoxicity of thiourea derivatives was assessed in HepG2 and HepG2.2.15 cells by MTT assay whereas their genotoxicity was measured by Ames II test. The acute toxicity study was carried out in the Sprague-Dawley rats by observing the following parameters: mortality, clinical symptoms, hematological parameters, urine and changes in animal body & organs weight, gross necropsy and histopathology. DSA-00, DSA-02, and DSA-09 were non-cytotoxic even at the **320μM** concentration with respective CC_50_ values of **329.6, 323.5,** and **349.7 μM**. The Ames II test revealed that these molecules were non-mutagenic at a ∼1M concentration. The acute toxicity studies revealed LD_50_ values belong from the moderate range of toxicity of DSA-00, DSA-02 and DSA-09. Importantly, there were no abnormal change in body weight and organs weight. Moreover, no abnormal clinical signs such as hematological parameter, urinalysis, gross necropsy and or histopathological in any of the animals after receiving oral doses of thiourea derivatives. Thiourea derivatives **DSA-00**, **DSA-02**, and **DSA-09** appear to have a moderate range of acute toxicity at dosages used and thus, appear to be generally safe.

**Graphical Abstract.**
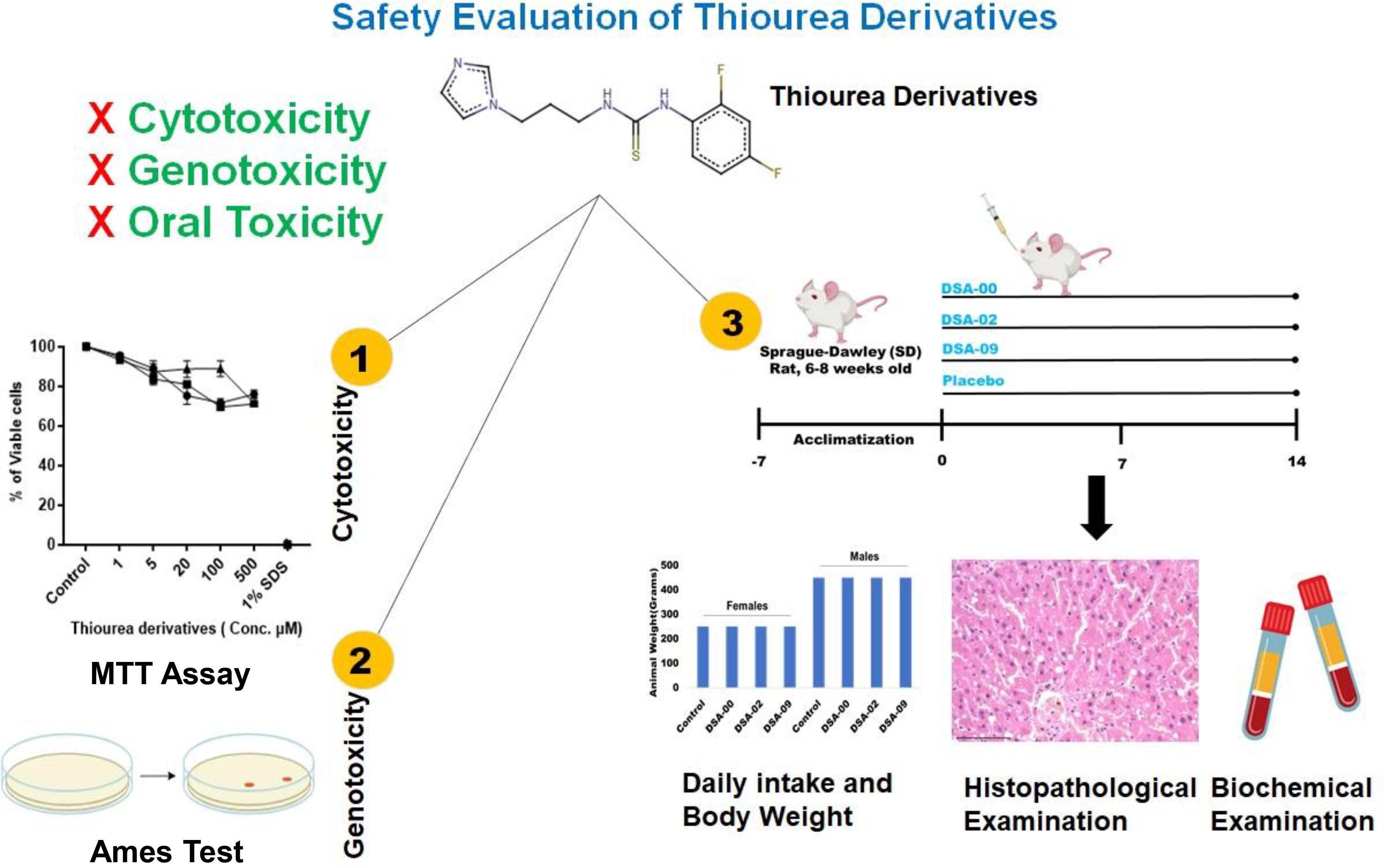
Illustration of the study design and the findings for each aspect of the study: in-vitro cytotoxicity (1), genotoxicity by the Ames test (2), and oral toxicity in rats (3).

## 1. Introduction

Organosulfur chemical compounds known as thiourea derivatives have received a lot of attention due to their pharmacological properties, such as anticancer [1], anti-bacterial [2], antiviral [3], anti-fungal [4,5], anti-diabetic [6], anti-malarial [7] and antinociceptive. [8] One of the most effective ways to boost a chemicals biopotency, bioavailability, and lipophilicity is to incorporate a halogen functional group (s) into the thiourea derivatives. Moreover, p-acetamido-benzaldehyde thiosemicarbazone (also known as thiacetazone), which is used for the treatment of *Mycobacterium tuberculosis* (TB), is also used along with thiourea derivatives.[9] The effectiveness of a thiourea derivative against *Mycobacterium tuberculosis* was also improved by an oxygen-containing side chain. [10] Although currently used antiviral and antibiotic drugs are safe but repeated treatment provides resistance to bacteria, fungus, and viruses due to their high rate of mutation, which has also become a widespread problem. Therefore, better and safer drugs are needed. To find out new drugs usually molecular docking as well as compound libraries screening have been used to identify novel pharmaceuticals [11]; Singh et al., 2023 communicated). In addition, a thiourea derivative known as N-(4-methyl-2-thiazolyl)-N’-phenylthiourea (MTPT) has shown anti-viral activity against the hepatitis B virus (HBV). In a study, MTPT was tested for its anti-viral activity against HBV in-vitro. The results showed that MTPT effectively inhibited the replication of HBV in HepG2.2.15 cells, which are a commonly used cell line for studying HBV infection.[12,13] The novel identified drugs have safety issues as well as adverse effects on the health of patients. Therefore, pharmacokinetics and pharmacodynamics, as well as toxicological evaluation of newly identified drugs, are essential.

Here, we evaluated the toxicological properties of three thiourea derivatives (DSA-00, DSA- 02, and DSA-09), which recently identified as new antivirals, that suppressed HBV infection (Singh et al., 2023 communicated). The mode of action of these derivatives, masks the effect of HBx proteins as results suppression of HBV replication and restricts HBx mediated heptocarcinogenesis (Kumar et al., 2023 communicated). Further, the pharmacokinetically evaluation of thiourea derivatives using both *in-silico* and *in-vivo* results suggested that, these thiourea derivatives to be good candidate drugs. These results necessitated the need of a safety assessment of thiourea derivatives. Therefore, the present study was designed to evaluate the toxicological properties of thiourea derivatives, such as in-vitro cytotoxicity, mutagenicity, and acute animal toxicity. The new thiourea derivatives are non-cytotoxic and non-mutagens at high concentrations. According to **CFR toxicity category**, these derivatives are fall under the category III of animal toxicity or moderate range of toxicity.

## 2. Materials and methods

### 2.1 Test substances

Thiourea derivatives (Figure 1) were dissolved in dimethyl sulfoxide (DMSO) and water 10mg/mL concentration and stored at -20°C or -80°C until used.

**FIGURE 1.**
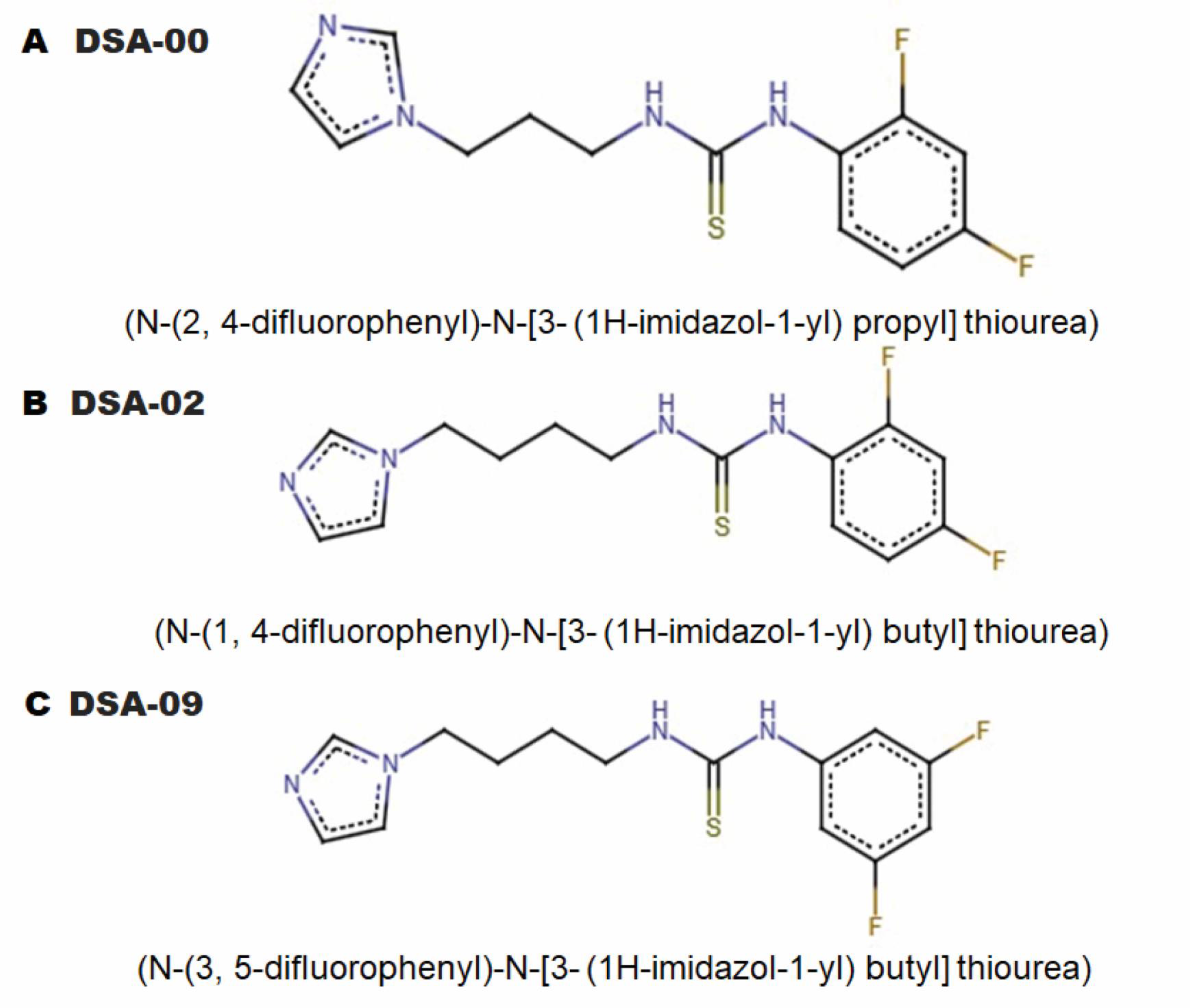
Sketch of chemical structure of thiourea derivatives. The chemical structures of thiourea derivatives (A= DSA-00, B= DSA-02, and C = DSA-09).

### 2.2 *In-vitro* cytotoxicity assay of thiourea derivatives

Thiazolyl Blue Tetrazolium Bromide powder (MTT, Cat no. TC191, HiMedia, India) was solubilized in phosphate buffer saline at 5 mg/mL concentration mixed by vertexing or sonication and filtered with 0.22µM Syringe filter (GSWP02500, Millipore). Solution aliquots were stored at 4°C for a week. Details protocol in supporting information.

### 2.3 Ames II test

There is an initial phase used to evaluate the dose range, and second phase used to determine mutagenicity in the Ames II test. The dose range-finding phase involves different concentrations of the Thiourea derivatives in triplicate for each strain, with and without S9 (Rat liver Extract) [14], more detail the strains used in the Ames test.[15] Detail’s protocol in supporting information.

### 2.4 Animal use and care

The study was carried out in accordance with the Organization for Economic Co-operation and Development (OECD) test guideline No. 425 (Up-Down procedure).[16] Healthy 6–8-week-old male and female Sprague Dawley (SD) rats were purchased from our animal house of Institute of Liver and Biliary Sciences, New Delhi, India after approved by Institution Animal Ethics Committee (IAEC/ILBS/20/08, 02/08/2020) for all the procedure applied in this research work. Animal were quarantined for a week then randomly assigned into five groups (5 animal each group), animals were housed in polypropylene cage at recommended standard conditions temperature (24 ± 2°C), humidity (55 ± 5%) and light/dark cycle (12 h light/ 12h dark with ad libitum food and water supply.

### 2.5 Single-oral dosage toxicity

#### 2.5.1 Single-oral dosage toxicity

For a single oral dosage toxicity examination. Rats were given vehicle (saline) or categorized dosages of thiourea derivatives (500, 1000, 2000, 3000 and 4500mg/kg) via oral gavage. After administration, animals were monitored for mortality and clinical symptoms every hour for 6 h and then daily for 14 days.

#### 2.5.2 Repeated oral dosage toxicity

For that, animals were treated with five dosages of thiourea derivatives (500, 1000, 2000, 3000 and 4500 mg/kg) via oral gavage once a day for 14 days while vehicle group animals were dosed with saline. Rats were regularly monitored for mortality and clinical symptoms throughout the experiment and animal bodyweight were also measured at 0, 7 and 14 days after post-treatment.

#### 2.5.3 Collection of blood and tissue samples

At the end of the 15 days of experiment, the Rats in each group were anaesthetized with pentobarbital sodium (60mg/kg-bw) and blood was collected through retro-orbital plexus into EDTA and non-EDTA tube for serum and other hematological analysis. For serum isolation, blood tube was left at room temperature for 30 minutes then centrifuged at 3000 RPM for 20 minutes and serum was collected into fresh Eppendorf tube and stored at -80oC till further process. Animals were sacrificed by using Carbon dioxide Chamber for 20 minutes continuous exposure of CO_2_ and organs were collected and stored into 10% formalin for the histopathological analysis further some portion of organs was snap freeze in to liquid nitrogen and stored at -80oC for further process. During animals sacrifices each body and organs weight was recorded and related organ to body weight was calculated using this formula. Relative organ weight [gm/100g body weight]

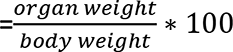

#### 2.5.4 Serum biochemistry

Blood serum used for measured the Alanine aminotransferase (ALT), Aspartate aminotransferase (AST), g-glutamyl transferase (GGT), Total protein (TP), albumin (ALB), globulin (G), albumin to globulin (A/G), total bilirubin (TBIL) Urea, urea nitrogen, and creatinine (CREA) were estimated using standard laboratory kits (ERBA), as per manufacturer’s instructions.

### 2.6 Histopathological Examination

For the histopathological examination, tissue samples were fixed into 10% formalin at room temperature for 24 h and paraffin blocks were prepared. 5µm thin sections were cut with the help of microtome and subjected to hematoxylin and eosin (H&E) stain according to standard protocol. Briefly, Tissue section were deparaffinized, rehydrated, dehydrated stained with Mayer’s Hematoxylin for 5–8 min, washed, and counterstained with Eosin Y. The stained slide was washed, dehydrated, mounted in DPX, and visualized under the light microscope.

### 2.7 Statistical analysis

Statistical tests were done using SPSS v25 and Graph Pad Prism a v8. Two-group comparisons were performed using univariate two-tailed student’s t-test analysis. Spearmen correlation analysis was performed, and R^2^>0.99 was considered statistically significant correlation.

## 3. Results

### 3.1 The cytotoxicity effect of thiourea derivatives

Our previous reports have suggested that, derivatives of thiourea did not showed any toxic effect in in-vitro. These thiourea derivatives viz. DSA-00, DSA-02, and DSA-09 have the antiviral potential to suppressed HBV Replication as comparable to currently used nucleosides analogous like Entecavir and Tenofovir (Singh et al., 2023 communication). Therefore, the need for testing cytotoxicity arises from the potential risks associated with these derivatives that can cause damage or death to cells in living organisms. However, there are no standard cytotoxicity test procedures, and each of the ones that do exist have their own set of drawbacks. Therefore, in-vitro cytotoxicity, which is commonly evaluated by the MTT assay, is one of the most important criteria in assessing toxicity biology. This method is widely accepted and continuously improved for modern cell biology.

Thus, the hepatoma cells were treated with concentration range of thiourea derivatives (1-500μM). Our observation revealed that DSA-00, DSA-02, and DSA-09 were safety vs non-cytotoxic at 320µM concentration (after calculation of CC_50_ value) *in-vitro* studies (Figure 2C-D), after conducting 24 h treatments with these drugs, the percentage of viable cells was calculated to be approximately 90-100% (Figure 2A). In both the HepG2.2.15 and HepG2- NTCP cell lines, the maximum number of cells that died was approximately 1-10%. These results were compared with cells treated with 1% Sodium dodecyl sulfate (SDS), which served as a positive control and resulted in 100% cell death (Figure 2B). It was observed that these thiourea derivatives were unable to cause cell death under *in-vitro* conditions. Therefore, it can be concluded that thiourea derivatives are safer even at higher concentrations (320µM).

**FIGURE 2.**
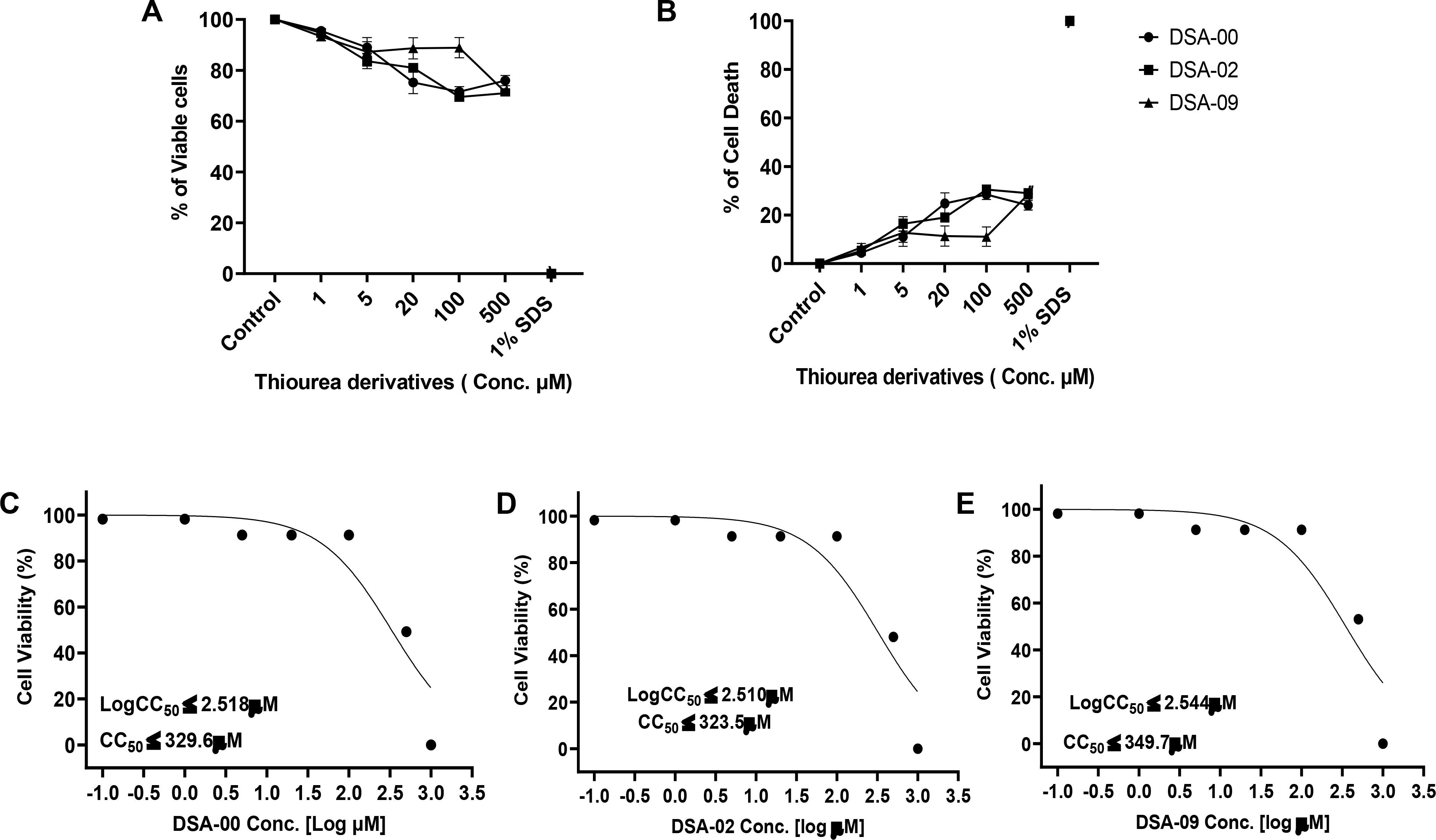
*In-vitro* cytotoxicity in the presence of DSA-00 and its two derivatives. HepG2- NTCP and HepG2.2.15, 2x10^4^ cells seeded in 96 well plates. After 24 h treated with 1 to 500 µM concentration of DSA-00 and its derivatives compounds for 24 hrs. Further calculated cells viability (A) and cell deaths(B) by MTT assay as per mentioned in methods in percentage. The Cells viability and cell deaths Each experiment was performed three times and in triplicate. The 50% cell death concentrations of DSA-00(C), DSA-02(D), and DSA-09(E). All experiments were performed thrice and in triplicate. Statistical significance was calculated using Student’s t test. *, p < 0.05; **, p < 0.001; ns=non-significant.

### 3.2 The mutagenic effect of thiourea derivatives

The mutagenicity of thiourea derivatives was assessed using the Ames II test, which is a widely accepted method for determining the mutagenic potential of chemical compounds. The test uses auxotrophic strains of Salmonella typhimurium that are unable to synthesize histidine or tryptophan due to point mutations in their genes. If a compound is mutagenic, it will cause a reversal of these mutations and allow the bacteria to grow on histidine- or tryptophan-deficient media.

In our study, we treated the TA98 and TA Mix (including TA7001-TA7006) bacterial strains with different concentrations (∼0.001-1M) of DSA-00, DSA-02, and DSA-09, with and without the presence of S9 (Rat Liver Extract), which is a metabolic activation system. We observed that the number of revertant colonies did not increase by more than 2 times compared to the negative control groups, indicating that the thiourea derivatives did not exhibit mutagenic activity. The TA98 bacterial strain was treated with 4-Nitroquinoline-N-Oxide (4-NQO, 50 μg/mL) and 2-Nitrofluorence (2-NF, 25 μg/mL) in the absence of S9 as a positive control, while the TA-mix was treated with 2-aminofluorene (2-AF, 125 μg/mL) in the presence of S9 as a positive control. These positive control compounds are known mutagens and have been shown to cause an increase in revertant colonies in the Ames II test (Table 1).

**Table 1.**
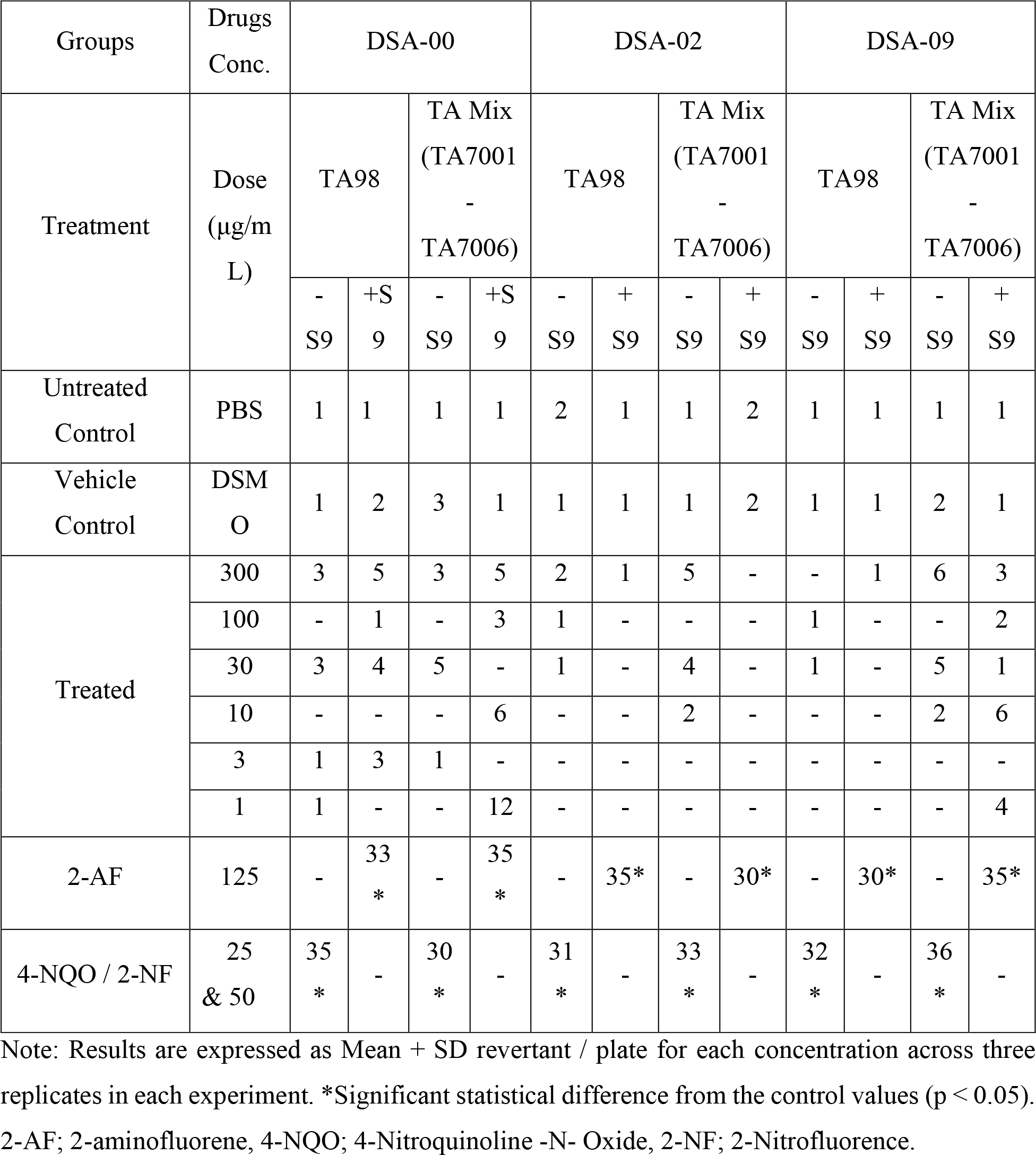
Ames test for DSA-00 and its derivatives DSA-02 & DSA-09

| Groups | Drugs Conc. | DSA-00 |  |  |  | DSA-02 |  |  |  | DSA-09 |  |  |  |
| --- | --- | --- | --- | --- | --- | --- | --- | --- | --- | --- | --- | --- | --- |
| Treatment | Dose (µg/mL) | TA98 |  | TA Mix (TA7001 - TA7006) |  | TA98 |  | TA Mix (TA7001 - TA7006) |  | TA98 |  | TA Mix (TA7001 - TA7006) |  |
|  |  | - S9 | +S9 | - S9 | +S9 | - S9 | + S9 | - S9 | + S9 | - S9 | + S9 | - S9 | + S9 |
| Untreated Control | PBS | 1 | 1 | 1 | 1 | 2 | 1 | 1 | 2 | 1 | 1 | 1 | 1 |
| Vehicle Control | DSMO | 1 | 2 | 3 | 1 | 1 | 1 | 1 | 2 | 1 | 1 | 2 | 1 |
| Treated | 300 | 3 | 5 | 3 | 5 | 2 | 1 | 5 | - | - | 1 | 6 | 3 |
|  | 100 | - | 1 | - | 3 | 1 | - | - | - | 1 | - | - | 2 |
|  | 30 | 3 | 4 | 5 | - | 1 | - | 4 | - | 1 | - | 5 | 1 |
|  | 10 | - | - | - | 6 | - | - | 2 | - | - | - | 2 | 6 |
|  | 3 | 1 | 3 | 1 | - | - | - | - | - | - | - | - | - |
|  | 1 | 1 | - | - | 12 | - | - | - | - | - | - | - | 4 |
| 2-AF | 125 | - | 33* | - | 35* | - | 35* | - | 30* | - | 30* | - | 35* |
| 4-NQO / 2-NF | 25 & 50 | 35* | - | 30* | - | 31* | - | 33* | - | 32* | - | 36* | - |

Note: Results are expressed as Mean + SD revertant / plate for each concentration across three replicates in each experiment. \*Significant statistical difference from the control values ( $p < 0.05$ ). 2-AF; 2-aminofluorene, 4-NQO; 4-Nitroquinoline -N- Oxide, 2-NF; 2-Nitrofluorence.

### 3.3 Effect of thiourea derivatives in single oral dose

Based on the above cytotoxicity and mutagenicity results, we next performed single-dose acute and 14-day repeated dose toxicity of thiourea derivatives (DSA-00, DSA-02, and DSA- 09) were as per the OECD 425 (Up-Down technique) guideline (16).The rats received various derivative doses and were observed for mortality and morphological, biochemical, clinical, and histological alterations during the 14-day experiments.

In the Up-Down procedure, all rats were administered 2000 mg/kg-bw of thiourea derivatives gastro intestinally (orally), and all rats survived during the 14 days of experiment. Therefore, 3000 mg/kg-bw was the dose for the second level administered, where two to three rats died during 14 days of experiments. Moreover, 4500 mg/kg-bw was the dose for the third level, and all rats died within two to three days. These observations suggested that the LD50 values for DSA-00, DSA-02, and DSA-09 were 2998.47, 3235.93, and 3235.93 mg/kg-bw, respectively (Figure 3, and Table 2). In either male or female rats, any dose studied did not significantly alter clinical symptoms, or macroscopic necropsy examination of any organs at postmortem (data not shown). Therefore, further evaluation of the 14-day repeated oral toxicity of these thiourea derivatives.

**FIGURE 3.**
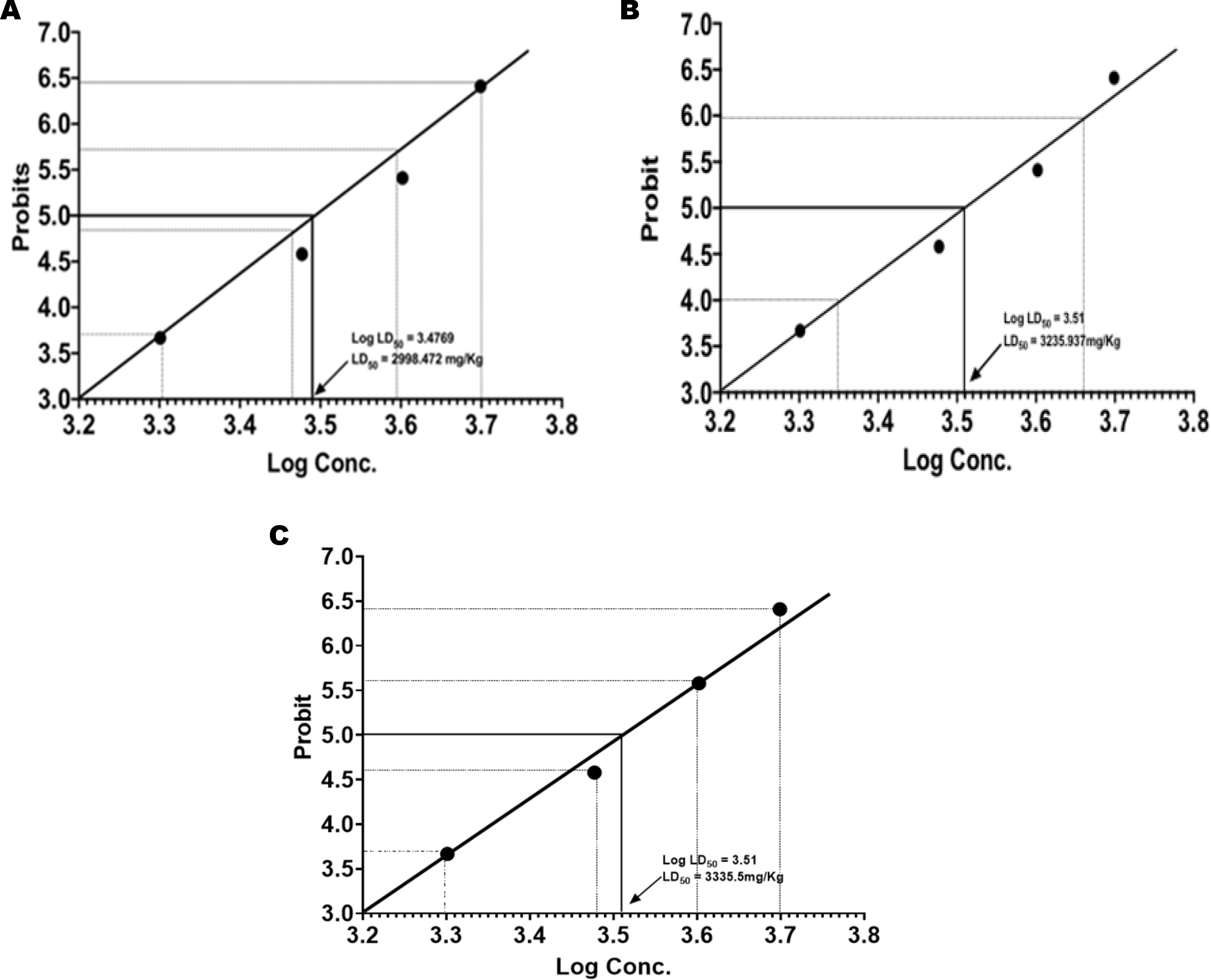
Determination of LD_50_ of thiourea derivatives. Thiourea derivatives were administered orally to randomly divided male and female rats weighing between 265g and 425g respectively. The concentration ranged from 500–4500 mg/kg-bw. Each group received a dose of 500, 1000, 2000, 3000, and 4500 mg/kg body weight in a volume given orally at a rate of 10 ml/kg body weight in a single dose. According to Miller and Tainter, the LD50 of DSA-00 (A), DSA-02 (B), and DSA-09 (C) were determined.

**Table 2.** Mortality after giving Rat DSA-00 via gavage and the related survival times.

| Dose<br>(mg/kg-bw) | Mortality |  |  | Survival Times (Days) |  |  |
| --- | --- | --- | --- | --- | --- | --- |
|  | DSA-00 | DSA-02 | DSA-09 | DSA-00 | DSA-02 | DSA-09 |
| 4500 | 2/3 | 3/3 | 3/3 | 2, & 2.5 | 1, 3, & 5 | 0.5, 3, & 4 |
| 3000 | 2/6 | 1/6 | 3/6 | 2 & 5 | 2.5 | 1.5, 3, & 5 |
| 2000 | 0/3 | 0/3 | 0/3 | >14 | >14 | >14 |
| 1500 | 0/3 | 0/3 | 0/3 | >14 | >14 | >14 |
| 1000 | 0/3 | 0/3 | 0/3 | >14 | >14 | >14 |
| 500 | 0/3 | 0/3 | 0/3 | >14 | >14 | >14 |

Note: Groups(n=3) based on dose (milligrams/kilograms-body weight), represented dead/live animal per sections in mortality and survival time.

### 3.4 Repeated oral dose toxicity of thiourea derivatives

Repeated doses of thiourea derivatives were administered over the course of 14 days for the evaluation of repeated oral dose toxicity; no deaths were detected within 4 h of continuous observations or within 24 h. Additionally, DSA-00, DSA-02, and DSA-09 were administered for 14 days; no adverse effects were observed in groups 500, 1000, 2000, and 3000, while in the group of 4500, two mortalities were observed between 3 and 14 days. The fur, skin, eyes, and nose, or morphological features, appeared normal. No aberrant behaviors, such as salivation, diarrhea, or lethargy, were noticed in the treated groups after 7 and 14 days as compared with 0 days. However, substantial drowsiness was noticed 10–15 minutes after the DSA-00 doses of 4500, 3000, 2000, 1000, and 500 mg/kg-bw were administered. That substantial drowsiness lasted for 40–50 minutes until it started to decrease, and normal activity was resumed within a few minutes. Throughout the course of the experiments, neither male nor female rats displayed any additional adverse side effects. These results indicate that DSA- 00, DSA-02, and DSA-09 at doses of 1000, 2000, and 3000 mg/kg-bw were safe because groups at 4500 mg/kg-bw observed mortality later during the period of experiments. Deaths were usually caused by the indigestion and deposition of thiourea derivatives in the stomach as well as their inability to digest in large amount of the molecules. Therefore, the optimized 3-level Up and Down procedure showed that DSA-00, DSA-02, and DSA-09-induced mortality were dose-independent. Therefore, based on these mortality data, we have calculated 50% death at the medium lethal dose (LD50) of thiourea derivatives.

### 3.5 Histopathological alterations and organ weights

We also checked and compared the macroscopic toxicities at 4500 mg/kg bw of DSA-00, DSA-02, and DSA-09 along with untreated (Placebo). No macroscopic toxicities were observed during in the 0, 7, or 14-day of observation periods. No obvious anomalies were found that may be ascribed to being treated with thiourea derivatives. The paraffins block of liver, kidney, ovary and testis, heart, spleen, ileum, and duodenum were prepared for histological examination. No discernible differences were observed in the control and thiourea derivatives groups (4500mg/kg) (Figure 4) and others groups results depicted in Figure S1- S8. Further, we checked the body weight, organ weight, organ to body weight ratio, feed intake, and water consumption after DSA-00, DSA-02, and DSA-09 administration for 14 days and observed that all these parameters were normal (Tables S1-S12)

**FIGURE 4.**
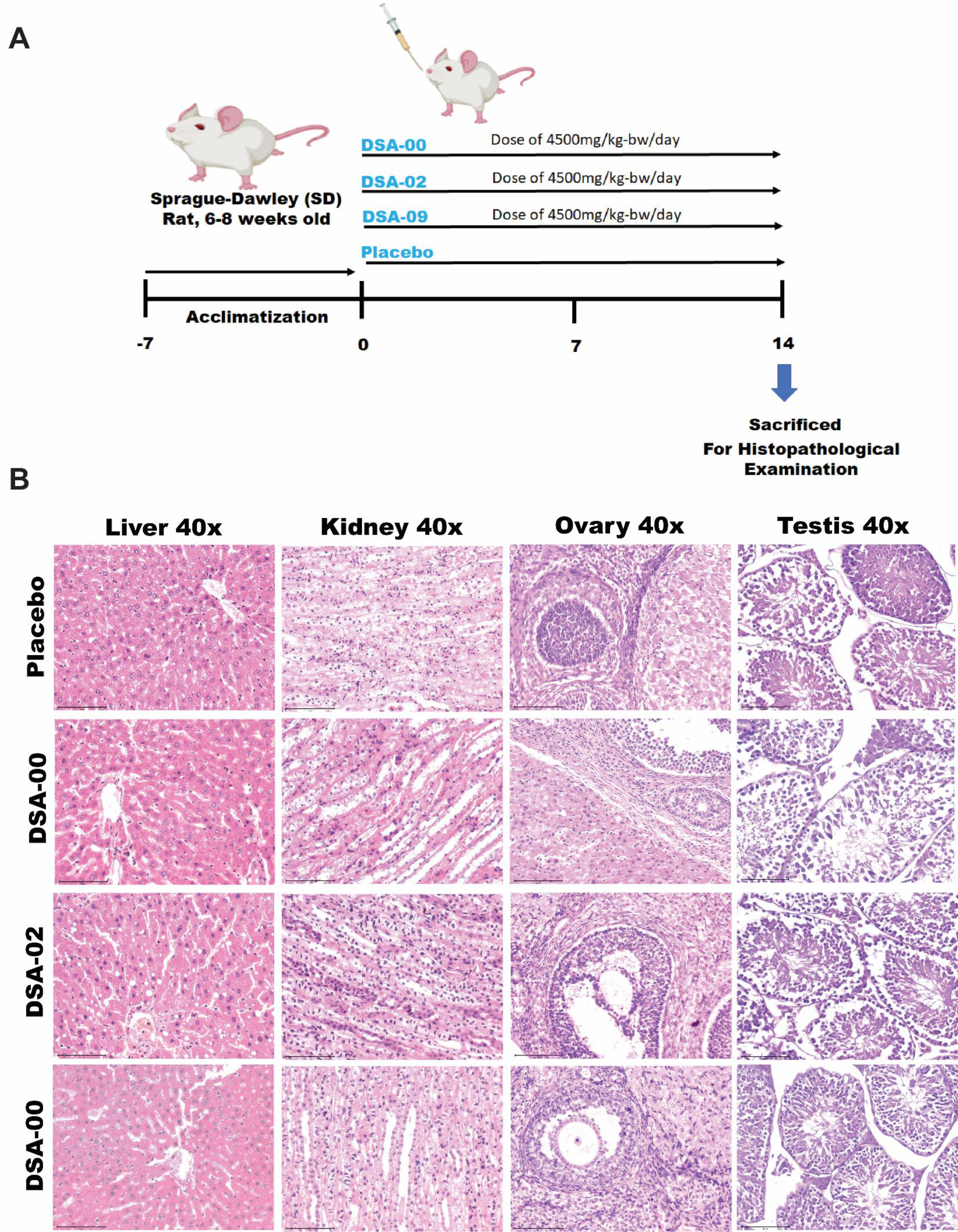
Illustration of Study flow(A,), the microphotographs (x40) showing the histological examination of the experimental rats that had been received 4500mg/kg-bw of thiourea Derivatives and compared with the control rats(B).

### 3.6 The evaluation of serum biochemistry

Next, we performed serum biochemical analysis of both males and female’s rats, specifically focusing on ALT, AST, GGT, and A/G levels. Both male and female rats showed no significant decrease in these parameters across treatments of DSA-00, DSA-02, and DSA-09. However, the ALT slightly increased in the treated with DSA-00 (34.2 ± 4.40) and DSA-02 (36.04 ± 1.73), when compared to the DSA-00 (32.2 ± 4.0) and DSA-02 (33.3 ± 1.86) treated females and the control group (29.68 ± 2.90). While the AST showed a little increase in the treated with DSA-02 (131.32 ± 4.05) as compared to the DSA-00 (121.88 ± 2.40) and DSA- 02 (107.32 ±12.99) treated females and control group (119.34 ± 3.92). In female subjects, the administration of DSA-00 resulted in an increase in TBIL levels as compared to the male subjects and females treated with DSA-02, and DSA-09, as well as no significant increase in palmatine levels compared to the females and control group. Furthermore, the study found no statistically significant effects on the remaining parameters (Table 3).

**Table 3.**
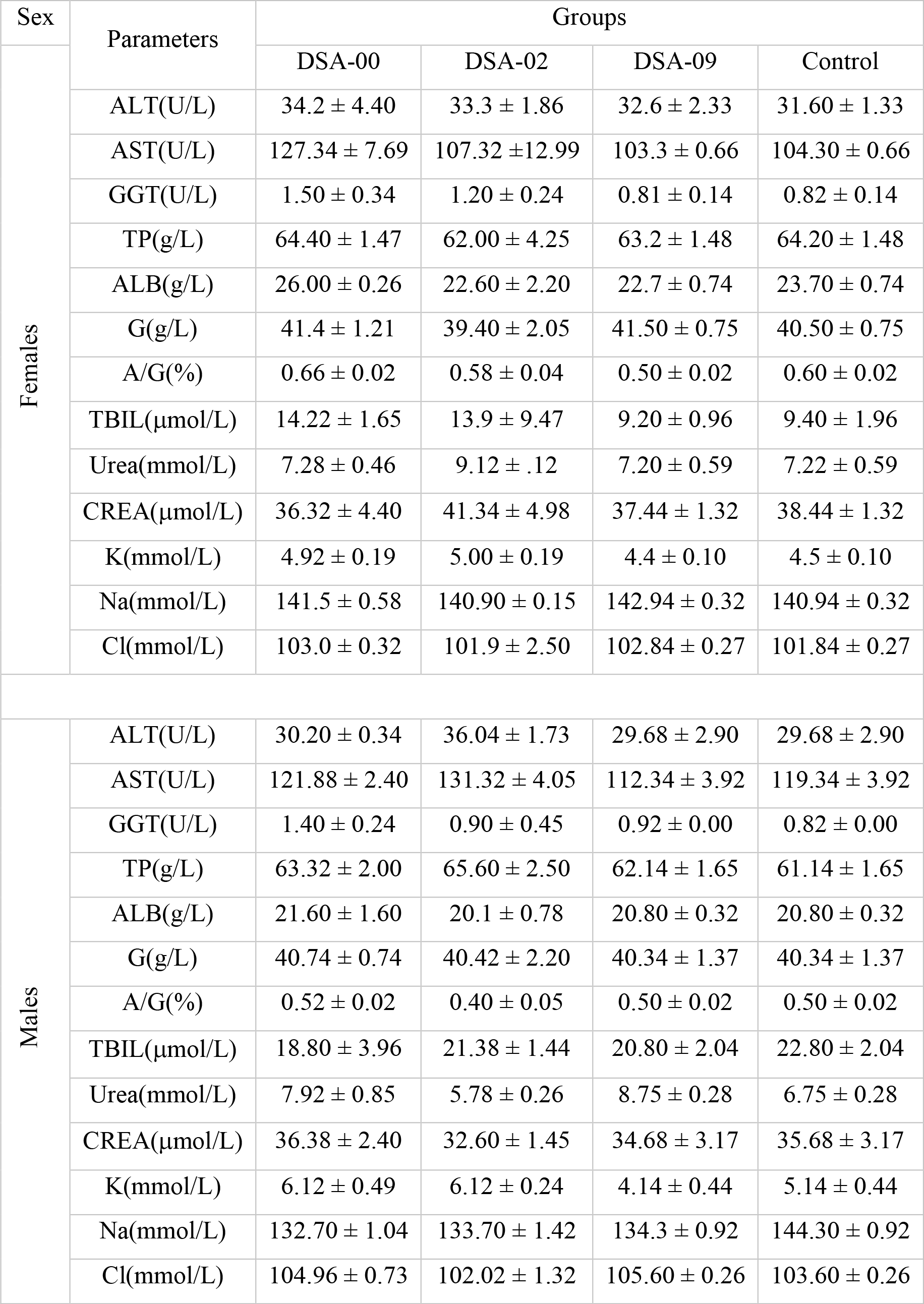
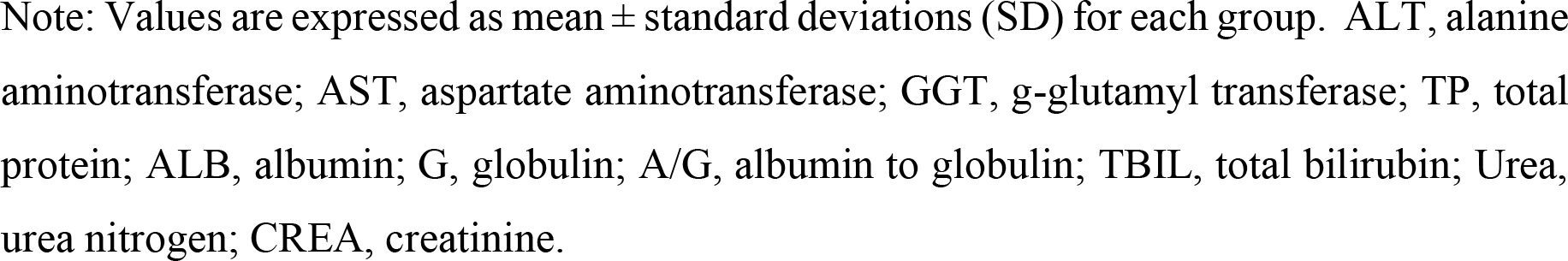
Hematological parameters of rats after thiourea derivatives treatment.

| Sex | Parameters | Groups |  |  |  |
| --- | --- | --- | --- | --- | --- |
| Females |  | DSA-00 | DSA-02 | DSA-09 | Control |
|  | ALT(U/L) | 34.2 ± 4.40 | 33.3 ± 1.86 | 32.6 ± 2.33 | 31.60 ± 1.33 |
|  | AST(U/L) | 127.34 ± 7.69 | 107.32 ± 12.99 | 103.3 ± 0.66 | 104.30 ± 0.66 |
|  | GGT(U/L) | 1.50 ± 0.34 | 1.20 ± 0.24 | 0.81 ± 0.14 | 0.82 ± 0.14 |
|  | TP(g/L) | 64.40 ± 1.47 | 62.00 ± 4.25 | 63.2 ± 1.48 | 64.20 ± 1.48 |
|  | ALB(g/L) | 26.00 ± 0.26 | 22.60 ± 2.20 | 22.7 ± 0.74 | 23.70 ± 0.74 |
|  | G(g/L) | 41.4 ± 1.21 | 39.40 ± 2.05 | 41.50 ± 0.75 | 40.50 ± 0.75 |
|  | A/G(%) | 0.66 ± 0.02 | 0.58 ± 0.04 | 0.50 ± 0.02 | 0.60 ± 0.02 |
|  | TBIL(μmol/L) | 14.22 ± 1.65 | 13.9 ± 9.47 | 9.20 ± 0.96 | 9.40 ± 1.96 |
|  | Urea(mmol/L) | 7.28 ± 0.46 | 9.12 ± .12 | 7.20 ± 0.59 | 7.22 ± 0.59 |
|  | CREA(μmol/L) | 36.32 ± 4.40 | 41.34 ± 4.98 | 37.44 ± 1.32 | 38.44 ± 1.32 |
|  | K(mmol/L) | 4.92 ± 0.19 | 5.00 ± 0.19 | 4.4 ± 0.10 | 4.5 ± 0.10 |
|  | Na(mmol/L) | 141.5 ± 0.58 | 140.90 ± 0.15 | 142.94 ± 0.32 | 140.94 ± 0.32 |
|  | Cl(mmol/L) | 103.0 ± 0.32 | 101.9 ± 2.50 | 102.84 ± 0.27 | 101.84 ± 0.27 |
| Males | ALT(U/L) | 30.20 ± 0.34 | 36.04 ± 1.73 | 29.68 ± 2.90 | 29.68 ± 2.90 |
|  | AST(U/L) | 121.88 ± 2.40 | 131.32 ± 4.05 | 112.34 ± 3.92 | 119.34 ± 3.92 |
|  | GGT(U/L) | 1.40 ± 0.24 | 0.90 ± 0.45 | 0.92 ± 0.00 | 0.82 ± 0.00 |
|  | TP(g/L) | 63.32 ± 2.00 | 65.60 ± 2.50 | 62.14 ± 1.65 | 61.14 ± 1.65 |
|  | ALB(g/L) | 21.60 ± 1.60 | 20.1 ± 0.78 | 20.80 ± 0.32 | 20.80 ± 0.32 |
|  | G(g/L) | 40.74 ± 0.74 | 40.42 ± 2.20 | 40.34 ± 1.37 | 40.34 ± 1.37 |
|  | A/G(%) | 0.52 ± 0.02 | 0.40 ± 0.05 | 0.50 ± 0.02 | 0.50 ± 0.02 |
|  | TBIL(μmol/L) | 18.80 ± 3.96 | 21.38 ± 1.44 | 20.80 ± 2.04 | 22.80 ± 2.04 |
|  | Urea(mmol/L) | 7.92 ± 0.85 | 5.78 ± 0.26 | 8.75 ± 0.28 | 6.75 ± 0.28 |
|  | CREA(μmol/L) | 36.38 ± 2.40 | 32.60 ± 1.45 | 34.68 ± 3.17 | 35.68 ± 3.17 |
|  | K(mmol/L) | 6.12 ± 0.49 | 6.12 ± 0.24 | 4.14 ± 0.44 | 5.14 ± 0.44 |
|  | Na(mmol/L) | 132.70 ± 1.04 | 133.70 ± 1.42 | 134.3 ± 0.92 | 144.30 ± 0.92 |
|  | Cl(mmol/L) | 104.96 ± 0.73 | 102.02 ± 1.32 | 105.60 ± 0.26 | 103.60 ± 0.26 |

Note: Values are expressed as mean ± standard deviations (SD) for each group. ALT, alanine aminotransferase; AST, aspartate aminotransferase; GGT, g-glutamyl transferase; TP, total protein; ALB, albumin; G, globulin; A/G, albumin to globulin; TBIL, total bilirubin; Urea, urea nitrogen; CREA, creatinine.

## 4. Discussion

The development of new drugs and enhancing the therapeutic potential of already-existing drugs both rely heavily on toxicological screening. [17,18] Regulatory bodies such as the OECD and the US Food and Drug Administration (FDA) assert that testing novel drugs in animals for pharmacological functions and potential toxicity is crucial. The FDA now regulates the harmful effects of chemicals, food additives, medications, and other substances in the twenty-first century.[16,19]

Our study aimed to investigate the cytotoxicity, mutagenicity, and acute toxicity of thiourea derivatives, DSA-00, DSA-02, and DSA-09, as potential candidates for antiviral therapy for chronic hepatitis B (CHB) infection (Singh et al., 2023 communicated). Previous reports suggest that thiourea derivatives have low cytotoxicity, but their cytotoxicity levels can differ depending on the assay, cell line, and experimental system utilized in their studies. The cytotoxicity of numerous new thiourea derivatives have been investigated in HeLa, MCF- 7, and A549 cancer cell lines and found that none of these exhibited a cytotoxic effect.[20] We also assessed the cytotoxicity of DSA-00, DSA-02, and DSA-09 on both HepG2.2.15 cells and HepG2-NTCP cells and found that even at higher concentrations (320µM), none of these derivatives cause significant cell death. These findings provide a basis for further investigations into the potential therapeutic applications of thiourea derivatives in treating diseases such as chronic hepatitis B infection.

There have been several studies that have evaluated the mutagenic potential of thiourea derivatives and identified some non-mutagenic compounds. For example, a study evaluated the genotoxic and mutagenic effects of seven thiourea derivatives using the Ames test and the comet assay. The results of this study showed that six out of seven tested compounds were non-mutagenic and non-genotoxic.[21] Another study evaluated the mutagenic potential of 10 thiourea derivatives using the Ames test and the chromosomal aberration assay. The results of this study showed that eight out of 10 tested compounds were non-mutagenic and non- genotoxic.[22] It is important to note, however, that the safety of any compound must be evaluated on a case-by-case basis, as the potential for mutagenicity may vary depending on the specific compound, the dose, the duration of exposure, and other factors. Therefore, it is important to conduct thorough safety evaluations of thiourea derivatives before considering their use in humans. Furthermore, our study evaluated the mutagenicity of the thiourea derivatives using the Ames II test, which showed no evidence of mutagenic effects either with or without S9 (rat liver extract). The thiourea derivatives, including DSA-00, DSA-02, and DSA-09, were found to be superior to all currently available drugs at a concentration of 300 µg. It is worth noting that the ETV-treated WT, Parp1-/-, Rad18-/-, and Brca1-/- cells showed higher H2AX accumulation in their nuclei, indicating enhanced DNA damage and ETV hypersensitivity.[23] In conclusion, our results indicate that the thiourea derivatives DSA-00, DSA-02, and DSA-09 did not exhibit mutagenic activity, even at higher concentrations. Therefore, these compounds may not be classified as mutagens and may have a low risk of causing genetic mutations.

The acute toxicity of a thiourea derivative can be evaluated by determining its LD50 value, which is the dose at which 50% of test animals die within a specified time period. However, the LD50 value can vary depending on the animal species, the route of administration, and the specific compound being tested. There have been some studies that have evaluated the acute toxicity of certain thiourea derivatives. A study evaluated the acute toxicity of a new thiourea derivative in rats. The results of this study showed that the LD50 value of the compound was greater than 5000 mg/kg in rats.[24–26] Similarly, another study evaluated the acute toxicity of a thiourea derivative in mice. The results of this study showed that the LD50 value of the compound was greater than 5000 mg/kg, which suggests that the compound has low acute toxicity in mice. It should be emphasized that the acute toxicity of thiourea derivatives can be influenced by various factors, such as the specific compound being tested and the experimental conditions used in the toxicity evaluation. Therefore, it is important to conduct thorough toxicity evaluations of thiourea derivatives before considering their use in humans. After the thiourea derivatives successfully passed the cytotoxicity and mutagenicity tests, the molecules were further evaluated for acute animal toxicity and LD50 values according to Miller and Tainter (1944).[27–29] The non-lethal toxicity was evaluated using the Up-down procedure from OECD guideline 425. For thiourea derivatives, we found the liver weight and the liver organ-to-body weight ratio did not significantly alter in the 500, 1000, 2000, 3000, and 4500 mg/kg-bw treatment groups compared to the control group. Male and female rats treated with DSA-00, DSA-02, and DSA-09 did not exhibit any discernible toxicity compared to the control group in histopathological results (Figure 4). Animals that received 4500 mg/kg bw of these compounds did not show any behavioral changes until 14 days, which is much higher than the level of antivirals studied in the past. Nevertheless, a detailed investigation is required to determine the long-term effects of thiourea derivatives on experimental animals.

## 5. Conclusion

The development of new drugs and enhancing the therapeutic potential of already-existing drugs both rely heavily on toxicological screening. Regulatory bodies such as the OECD and the FDA assert that testing novel drugs in animals for pharmacological functions and potential toxicity is crucial. Our study aimed to investigate the cytotoxicity, mutagenicity, and acute toxicity of thiourea derivatives, DSA-00, DSA-02, and DSA-09, as potential candidates for antiviral therapy for CHB infection. The results of this study suggest that thiourea derivatives, DSA-00, DSA-02, and DSA-09, are safe for use in in-vitro studies at higher concentrations (320µM), as they did not cause significant cell death. Our study evaluated the mutagenicity of the thiourea derivatives using the Ames II test, which showed no evidence of mutagenic effects either with or without S9. Therefore, these compounds may not be classified as mutagens and may have a low risk of causing genetic mutations. After the thiourea derivatives successfully passed the cytotoxicity and mutagenicity tests, the molecules were further evaluated for acute animal toxicity and LD50 values. For thiourea derivatives, we found that the liver weight and the liver organ-to-body weight ratio did not significantly alter. It is important to note that while the studies have shown promising results in terms of safety, further research is needed to evaluate their efficacy and safety in humans before they can be used as drugs for treating chronic HBV infections and related diseases.

## Supporting information

https://drive.google.com/drive/u/0/

## Acknowledgements

This work was supported by grants to VK from the Department of Biotechnology (DBT), Government of India (Grant No. BT/PR30082/Med/29/1341/2018) and J.C. Bose National Fellowship (Grant No. SR/S2/JCB-80)/2012) of the Department of Science and Technology, Government of India, New Delhi. JK, and PT received Senior Research Fellowships respectively from the Council of Scientific and Industrial Research, University Grants Commission, New Delhi, and AKS received Senior Research Fellowship from Department of Biotechnology Government of India, New Delhi, New Delhi for the period of this study.

