## Supplementary material for "Safety Assessment and Evaluation of Novel Thiourea Derivatives as Antivirals": https://drive.google.com/drive/u/0/

**Method 1** Ames II test protocol.

**Table S1** Weight gain in rats during treatment with DSA-00.

**Table S2** Food intake during treatment with DSA-00.

**Table S3** Effect of DSA-00 on the weight of organs of rats treated for 14 days

**Table S4** Change in the organ to bodyweight ratio (g/100g bodyweight) in the rats treated for 14 days with DSA-00

**Table S5** Weight gain in rats during treatment with DSA-00.

**Table S6** Food intake during treatment with DSA-00.

**Table S7** Effect of DSA-00 on the weight of organs of rats treated for 14 days

**Table S8** Change in the organ to bodyweight ratio (g/100g bodyweight) in the rats treated for 14 days with DSA-00

**Table S9** Weight gain in rats during treatment with DSA-00.

**Table S10** Food intake during treatment with DSA-00.

**Table S11** Effect of DSA-00 on the weight of organs of rats treated for 14 days

**Table S12** Change in the organ to bodyweight ratio (g/100g bodyweight) in the rats treated for 14 days with DSA-00.

**FIGURE S1** The microphotographs (x40) of spleen at 1000, 2000, 3000, 4500 mg/kg-bw groups of thiourea derivatives treated and placebo males and females.

**FIGURE S2** The microphotographs (x40) of heart at 1000, 2000, 3000, 4500 mg/kg-bw groups of thiourea derivatives treated and placebo males and females.

**FIGURE S3** The microphotographs (x40) of kidney at 1000, 2000, 3000, 4500 mg/kg-bw groups of thiourea derivatives treated and placebo males and females.

**FIGURE S4** The microphotographs (x40) of Liver at 1000, 2000, 3000, 4500 mg/kg-bw groups of thiourea derivatives treated and placebo males and females.

**FIGURE S5** The microphotographs (x40) of Testis at 1000, 2000, 3000, 4500 mg/kg-bw groups of thiourea derivatives treated and placebo males.

**FIGURE S6** The microphotographs (x40) of Ovary at 1000, 2000, 3000, 4500 mg/kg-bw groups of thiourea derivatives treated and placebo females.

**FIGURE S7** The microphotographs (x40) of Ileum at 1000, 2000, 3000, 4500 mg/kg-bw groups of thiourea derivatives treated and placebo males and females.

**FIGURE S8** The microphotographs (x40) of Duodenum at 1000, 2000, 3000, 4500 mg/kg-bw groups of thiourea derivatives treated and placebo males and females.

**Method 1. In-vitro cytotoxicity and Cell viability measured by MTT**

Thiazolyl Blue Tetrazolium Bromide powder (MTT), (HiMedia, #TC191) was solubilized in phosphate buffer saline(PBS) at 5 mg/mL concentration mixed by vortexing or sonication and filtered with a 0.22uM Syringe filter (GSWP02500, Millipore). Solutions were aliquoted and stored at 4°C for a week.

20K cells (HepG2-NTCP and HepG2.2.15) were seeded in 96 well plates 9((167008, Nunc, Thermo) treated with DSA-00(IR-415) derivatives for 24 hrs. from 1 to 500µM concentration. On the second day, 50 µL of serum-free media and 50 µL of MTT solution was added to each well. After incubation for 3 hours at 37°C, 150 µL, DMSO was added to each well. Wrapped plate in foil and shaken on an orbital shaker for 15 minutes. Absorbance was recorded at OD=590 nm. Read plate within 30 minutes. Further calculated the cell cytotoxicity via taken mean of triplicate reading for each sample and subtracted with the culture medium background from assay readings. Finally calculated the percentage cytotoxicity with the followed equation, using corrected absorbance: % cytotoxicity = (100 x (control - sample)).[1-3]

**Method 2. Ames II test protocol**

**Test compounds Exposure;** The test compounds were exposed using a modified liquid fluctuation test in TAMix and TA98. Ames II Exposure media (Moltox, Boone, NC) was added to each well of a 24-well plate along with 0.050 ml of each overnight culture in the absence of S9 fraction. A 0.010 ml aliquot of each test substance was introduced. In tests using rat liver S9 fraction induced by Aroclor-1254 (Moltox, Boone, NC), the exposure medium aliquot was reduced to 0.152 ml to accommodate 0.038 ml of the S9 reagent. A final concentration of 4.5% S9 fraction was provided as a result. The S9 mixture included 102 mM $NaH_2PO_4$ buffer, 102 mM KCl, 8 mM $MgCl_2$, 5 mM glucose-6-phosphate, 4 mM nicotinamide adenine dinucleotide, and 30% S9 (Moltox, Boone, NC). The 24-well plates were shaken at 250 rpm while being incubated at 37°C for 90 min.

**Mutant Selection;** The 24-well plates were taken out of the incubator and placed on the platform of a robotics station after the 90-minute incubation period. The pipet dispensed an aliquot of 2.8 ml of histidine-deficient Ames II Reversion Indicator media (Moltox, Boone, NC) into each well of the 24-well plates containing chemically treated cultures. This successfully reduced any histidine that was still present in the exposure medium, halting the expansion of the auxotrophic colony. In the indicator medium that chooses for prototrophic reversion was gently mixed several times. The pipetting then distributed each well of a 24-well microliter plate in 50µl aliquots over 48 wells of a 384-well microliter plate. Each sample was effectively divided throughout 48 wells of the plate by transferring each column (four wells) of the 24-well plate into one half of a 384-well plate. As a result, one plate was utilized for each strain and copy. The 384-well microliter plates were incubated at 37°C for 48 hours after being wrapped in Ziploc plastic bags to avoid evaporation.

**Data Acquisition;** As the pH decreases (pKa ~ 5.2), a pH indicator dye in the Indicator Media turns yellow because catabolites accumulate from the metabolically active revertant cells that proliferate in the absence of histidine. In order to determine the frequency of reversion per duplicate per dose, the number of yellow positive wells out of a total of 48 wells was compared to the number of spontaneously revertant wells found in the solvent control sections. An SLT Spectra Image plate reader (Thermo Fisher, USA) was used to score each 48-well portion of the 384-well plates for the number of revertant wells (yellow), using optical density $(OD)_{492}$ nm as a reference wavelength and normalizing it to $OD_{623}$ nm. The SLT data Capture software digitized the optical density and sent the data to Microsoft Excel. In order to classify the data, summary tables were generated for each compound code.

There were no replicates used for the initial screen. When analyzing test compounds using single data points, the focus must be on dose dependency rather than single data points. A single, isolated data point that exceeds the predetermined baseline cutoff of "zero dose plus 1 SD" has limited significance and is ineligible for the designation of "positive" or "weak positive" for a chemical. A single data point, though. At the highest concentration examined, a 4-fold increase over baseline would signify the start of a dose-dependent response, hence this result was labelled as "possibly positive." Table II provides a summary of the guidelines used in the evaluation of compounds that underwent a single test.

**Method 3. Acute animal toxicity studies**

Acute toxicity study was conducted employing male and female in half rats in order to check whether DSA-00 and its derivatives produced any mortality and adverse reactions when given intragastrically at maximal feasible dosage in rats. Rats were randomly divided into 4 groups (control group, DSA-00, DSA-02, and DSA-09 treatment group) for each chemical compounds, consisting of fifteen animals (8 males and 7 females) in each group and three rats of the same gender from the same group were held in same box. The body weight range at the outlet of treatment was 420 ± 25g for males and 265 ± 20g for females. The compounds were administered orally (using a ball-tipped incubation steel needle fitted onto a graded disposable syringe) as a single bolus dose (5000mg/kg-bw), which is the maximum technically feasible dose in rats the maximum practicable concentration and volume that can be given intragastrically. Control rats received intragastrically administration of sterile saline (10ml/kg-bw) alone. Prior to dosing, rats were fasted overnight to eliminate feed from gastrointestinal tract. All animals were thoroughly observed after test compounds administration immediately for the onset of any toxic signs and once daily thereafter for 14 days of observation period to record any delayed toxic effects. Survival, feed intake (days 2 and 8), and body weight (days 0, 3, 7, 10, and 13) were monitored. On day 15, rats were anesthetized with pentobarbital sodium (60mg/kg-bw). The blood parameters were analyzed with the help of automatic biochemical analyzer, using standard ELISA and CLIA kits.

| **Sex** | **Animal No.** | **Initial Weight (g)** | **7-day (g)** | **14-day (g)** | **Total (g)** |
|---|---|---|---|---|---|
| **Female** | 1 | 265.00±15.00 | 295.00±25.00 | 330.00±15.00 | 70.00±5.00 |
| | 2 | 265.00±15.00 | 295.00±26.00 | 320.00±16.00 | 50.00±9.00 |
| | 3 | 265.00±15.00 | 285.00±27.00 | 327.00±17.00 | 67.00±7.00 |
| | 4 | 265.00±16.00 | 275.00±28.00 | 315.00±18.00 | 50.00±12.00 |
| | 5 | 265.00±14.00 | 28.00±29.00 | 337.00±19.00 | 62.00±8.00 |
| **Male** | 1 | 425.00±27.00 | 49.00±4.00 | 515.00±6.00 | 90.00±15.00 |
| | 2 | 425.00±23.00 | 49.00±41.00 | 517.00±61.00 | 92.00±13.00 |
| | 3 | 425.00±2.00 | 49.00±42.00 | 550.00±62.00 | 80.00±13.00 |
| | 4 | 425.00±21.00 | 49.00±43.00 | 518.00±63.00 | 93.00±16.00 |
| | 5 | 425.00+25.00 | 49.00+44.00 | 545.00±64.00 | 90.00±20.00 |

Table S1. Weight gain in rats during treatment with DSA-00.

| **Sex** | **Animal No.** | **In 7-day (g)** | **In 7-14-day (g)** | **Food Total (g)** |
|---|---|---|---|---|
| **Female** | 1 | 105.15 ±2.30 | 107.89 ±9.50 | 213.04 ±7.60 |
| | 2 | 106.70 ±3.60 | 140.40 ±18.50 | 247.1 ±15.36 |
| | 3 | 108.62 ±2.15 | 115.00 ±2.30 | 223.62 ±14.63 |
| | 4 | 105.73 ±5.90 | 111.15 ±16.70 | 216.88 ±17.56 |
| | 5 | 106.70 ±3.50 | 99.22±19.1 | 205.92±15.6 |

| | | | | |
|---|---|---|---|---|
| **Male** | 1 | 155.00 ±6.70 | 113.67±8.35 | 268.67±5.36 |
| | 2 | 141.51 ±15.20 | 117.89±11.2 | 258.41±12.52 |
| | 3 | 148.26 ±18.1 | 115.6±16.5 | 263.86±14.65 |
| | 4 | 131.15 ±14.2 | 114.63±15.36 | 245.78±13.25 |
| | 5 | 149.59±16.3 | 140.40±19.26 | 193.63±18.32 |

Table S2. Food intake during treatment with DSA-00.

| Sex | Dose (mg/kg-BW) | Body weight before sacrifice | Liver (g) | Kidney (g) | Spleen (g) | Testis/Ovary (g) |
|---|---|---|---|---|---|---|
| | 1 | 335.00±15.00 | 13.75±0.85 | 3.50±0.22 | 0.62±0.15 | 0.22±0.10 |
| | 2 | 325.00±16.00 | 13.60±0.95 | 3.15±0.25 | 0.71±0.15 | 0.21± 0.15 |
| **Female** | 3 | 327.00±17.00 | 12.72±0.70 | 3.20±0.30 | 0.68±0.13 | 0.24±0.12 |
| | 4 | 315.00±18.00 | 11.50±0.96 | 3.15±0.36 | 0.65±0.10 | 0.17±0.15 |
| | 5 | 337.00±19.00 | 13.60±0.89 | 3.00±0.75 | 0.64±0.16 | 0.19±0.11 |
| | 1 | 515.00±6.00 | 16.00±0.90 | 4.62±0.86 | 0.85±0.20 | 4.15±0.35 |
| | 2 | 517.00±61.00 | 15.15±0.98 | 4.23±0.79 | 0.90±0.22 | 4.1±0.33 |
| **Male** | 3 | 550.00±62.00 | 15.50±0.75 | 4.15±0.56 | 0.87±0.18 | 3.9±0.43 |
| | 4 | 518.00±63.00 | 17.10±0.85 | 4.36±0.81 | 0.82±0.17 | 4.0±0.30 |
| | 5 | 545.00±64.00 | 16.00±0.95 | 4.26±0.75 | 0.97±0.25 | 4.0±0.50 |

Table S3. Effect of DSA-00 on the weight of organs of rats treated for 14 days.

| Sex | Dose (mg/k-bw) | Bodyweight before sacrifice | Liver (g/100g) | Kidney (g/100g) | Spleen (g/100g) | Testis /Ovary (g/100g) |
|---|---|---|---|---|---|---|
| | 1 | 335.00±15.00 | 4.10±0.60 | 0.91±0.11 | 0.19±0.2 | 0.7±0.10 |
| | 2 | 325.00±16.00 | 4.18±0.75 | 0.97±0.90 | 0.22±0.200 | 0.6±0.10 |
| **Female** | 3 | 327.00±17.00 | 3.89±0.55 | 0.98±0.13 | 0.21±0.10 | 0.7±0.20 |
| | 4 | 315.00±18.00 | 3.65±0.53 | 1.00±0.80 | 0.21±0.40 | 0.5±0.10 |
| | 5 | 337.00±19.00 | 4.40±0.54 | 0.89±0.11 | 0.19±0.40 | 0.6±0.10 |
| | 1 | 515.00±6.00 | 3.11±0.47 | 0.9±0.10 | 0.17±0.30 | 0.9±0.60 |
| | 2 | 517.00±61.00 | 2.93±0.44 | 0.82±0.70 | 0.17±0.50 | 0.91±0.80 |
| **Male** | 3 | 555.00±62.00 | 2.98±.55 | 0.82±0.70 | 0.17±0.40 | 0.89±0.40 |
| | 4 | 518.00±63.00 | 3.3±.43 | 0.84±0.80 | 0.16±0.30 | 0.91±0.90 |
| | 5 | 545.00±64.00 | 2.94±.48 | 0.78±0.60 | 0.18±0.90 | 0.92±0.70 |

Table S4. Change in the organ to bodyweight ratio (g/100g bodyweight) in the rats treated for 14 days with DSA and its derivatives.

| Sex | Dose (mg/kg-BW) | Initial Weight (g) | 7-day (g) | 14-day (g) | Total (g) |
|---|---|---|---|---|---|
| **Female** | 1 | 265.00±15.00 | 290.00±25.00 | 335.00±15.00 | 70.00±5.00 |
| | 2 | 265.00±15.00 | 291.00±26.00 | 325.00±16.00 | 60.00±9.00 |
| | 3 | 265.00±15.00 | 280.00±27.00 | 327.00±17.00 | 62.00±7.00 |
| | 4 | 265.00±16.00 | 275.00±28.00 | 315.00±18.00 | 50.00±12.00 |
| | 5 | 265.00±14.00 | 28.00±29.00 | 337.00±19.00 | 62.00±8.00 |
| **Male** | 1 | 425.00±27.00 | 49.00±40.00 | 515.00±6.00 | 90.00±15.00 |
| | 2 | 425.00±23.00 | 49.00±41.00 | 517.00±61.00 | 92.00±13.00 |
| | 3 | 425.00±2.00 | 49.00±42.00 | 550.00±62.00 | 80.00±13.00 |
| | 4 | 425.00±21.00 | 49.00±43.00 | 518±63.00 | 93.00±16.00 |
| | 5 | 425.00±25.00 | 49.00±44.00 | 545.00±64.00 | 90.00±20.00 |

Table S5. Weight gain in rats during treatment with DSA-02**.**

| Sex | Animal No. | In 7-days (g) | In 14-days (g) | Total (g) |
|---|---|---|---|---|
| **Female** | 1 | 11.15 ±12.30 | 17.89 ±9.50 | 29.40 ±7.60 |
| | 2 | 86.70 ±8.60 | 14.4 ±18.50 | 19.74 ±15.36 |
| | 3 | 88.62 ±15.00 | 15.00 ±2.30 | 193.63 ±14.63 |
| | 4 | 85.73 ±18.90 | 11.15 ±16.70 | 186.88 ±17.56 |
| | 5 | 86.70 ±13.50 | 99.22±19.10 | 185.92±15.60 |
| **Male** | 1 | 15.00 ±6.70 | 113.67±8.35 | 218.67±5.36 |
| | 2 | 91.51 ±15.20 | 17.89±11.20 | 199.41±12.52 |
| | 3 | 98.26 ±18.10 | 115.6±16.50 | 213.86±14.65 |
| | 4 | 11.15 ±14.20 | 114.63±15.36 | 215.78±13.25 |
| | 5 | 89.59±16.30 | 14.40±19.26 | 193.63±18.32 |

Table S6. Food intake during treatment with DSA-02.

| Sex | Animal No. | Body weight before sacrifice | Liver (g) | Kidney (g) | Spleen (g) | Testis/Ovary (g) |
|---|---|---|---|---|---|---|
| **Female** | 1 | 335.00±15.00 | 13.75±0.85 | 3.50±0.22 | 0.62±0.15 | 0.22±0.10 |
| | 2 | 325.00±16.00 | 13.60±0.95 | 3.15±0.25 | 0.71±0.15 | 0.21± 0.15 |
| | 3 | 327.00±17.00 | 12.72±0.7 | 3.20±0.3 | 0.68±0.13 | 0.24±0.12 |
| | 4 | 315.00±18.00 | 11.50±0.96 | 3.15±0.36 | 0.65±0.10 | 0.17±0.15 |
| | 5 | 337.00±19.00 | 13.60±0.89 | 3.00±0.75 | 0.64±0.16 | 0.19±0.11 |
| **Male** | 1 | 515.00±6.00 | 16.00±0.9 | 4.62±0.86 | 0.85±0.20 | 4.15±0.35 |
| | 2 | 517.00±61.00 | 15.15±0.98 | 4.23±0.79 | 0.90±0.20 | 4.1±0.33 |
| | 3 | 550.00±62.00 | 15.50±0.75 | 4.15±0.56 | 0.87±0.18 | 3.9±0.43 |
| | 4 | 518.00±63.00 | 17.10±0.85 | 4.36±0.81 | 0.82±0.17 | 4.0±0.30 |
| | 5 | 545.00±64.00 | 16.00±0.95 | 4.26±0.75 | 0.97±0.25 | 4.0±0.50 |

Table S7. Effect of DSA-02 on the weight of organs of rats treated for 14 days.

| Sex | Dose (mg/k-bw) | Bodyweight before sacrifice | Liver (g/100g) | Kidney (g/100g) | Spleen (g/100g) | Testis /Ovary (g/100g) |
|---|---|---|---|---|---|---|
| **Female** | 1 | 335.00±15.00 | 4.10±0.60 | 0.91±0.11 | 0.19±0.20 | 0.7±0.10 |
| | 2 | 325.00±16.00 | 4.18±0.75 | 0.97±0.90 | 0.22±0.20 | 0.6±0.10 |
| | 3 | 327.00±17.00 | 3.89±0.55 | 0.98±0.13 | 0.21±0.10 | 0.7±0.20 |
| | 4 | 315.00±18.00 | 3.65±0.53 | 1.00±0.80 | 0.21±0.40 | 0.5±0.10 |
| | 5 | 337.00±19.00 | 4.40±0.54 | 0.89±0.11 | 0.19±0.40 | 0.6±0.10 |
| **Male** | 1 | 515.00±6.00 | 3.11±0.47 | 0.9±0.10 | 0.17±0.30 | 0.9±0.60 |
| | 2 | 517.00±61.00 | 2.93±0.44 | 0.82±0.70 | 0.17±0.50 | 0.91±0.80 |
| | 3 | 550.00±62.00 | 2.98±.55 | 0.82±0.70 | 0.17±0.40 | 0.89±0.40 |
| | 4 | 518.00±63.00 | 3.3±.43 | 0.84±0.80 | 0.16±0.30 | 0.91±0.90 |
| | 5 | 545.00±64.00 | 2.94±.48 | 0.78±0.60 | 0.18±0.90 | 0.92±0.70 |

Table S8. Change in the organ to bodyweight ratio (g/100g bodyweight) in the rats treated for 14 days with DSA and its derivatives.

| Sex | Animal No. | Initial Weight (g) | 7-day (g) | 14-day (g) | Total (g) |
|---|---|---|---|---|---|
| Female | 1 | 265.00+15.00 | 291.00+25.00 | 335.00+15.00 | 71.00+5.00 |
| | 2 | 265.00+15.00 | 296.00+26.00 | 325.00+16.00 | 66.00+9.00 |
| | 3 | 265.00+15.00 | 284.00+27.00 | 331.00+17.00 | 68.00+7.00 |
| | 4 | 265.00+16.00 | 270.00+28.00 | 310.00+18.00 | 40.00+12.00 |
| | 5 | 265.00+14.00 | 280.00+29.00 | 337.00+19.00 | 62.00+8.00 |
| Male | 1 | 425.00+27.00 | 490.00+14.00 | 515.00+26.00 | 90.00+15.00 |
| | 2 | 425.00+23.00 | 490.00+15.00 | 517.00+21.00 | 92.00+13.00 |
| | 3 | 425.00+2.00 | 490.00+13.00 | 550.00+22.00 | 80.00+13.00 |
| | 4 | 425.00+21.00 | 490.00+14.00 | 518.00+23.00 | 93.00+16.00 |
| | 5 | 425.00+25.00 | 490.00+14.00 | 545.00+24.00 | 90.00+12.00 |

Table S9. Weight gain in rats during treatment with DSA-09.

| Sex | Animal No. | 7-day (g) | 14-day (g) | Total (g) |
|---|---|---|---|---|
| Female | 1 | 11.15 +12.30 | 17.89 +9.50 | 29.40 +7.60 |
| | 2 | 86.70 +8.60 | 14.40 +18.50 | 19.74 +15.36 |
| | 3 | 88.62 +15.00 | 15.00 +2.30 | 193.63 +14.63 |
| | 4 | 85.73 +18.90 | 11.15 +16.70 | 186.88 +17.56 |
| | 5 | 86.70 +13.50 | 99.22+19.10 | 185.92+15.60 |
| Male | 1 | 15.00 +6.70 | 113.67+8.35 | 218.67+5.36 |
| | 2 | 91.51 +15.20 | 17.89+11.20 | 199.41+12.52 |
| | 3 | 98.26 +18.10 | 115.60+16.50 | 213.86+14.65 |
| | 4 | 11.15 +14.20 | 114.63+15.36 | 215.78+13.25 |
| | 5 | 89.59+16.30 | 14.40+19.26 | 193.63+18.32 |

Table S10. Food intake during treatment with DSA-09.

| Sex | Animal No. | Body weight before sacrifice | Liver (g) | Kidney (g) | Spleen (g) | Testis/Ovary (g) |
|---|---|---|---|---|---|---|
| **Female** | 1 | 335.00±15.00 | 13.75±0.85 | 3.50±0.22 | 0.62±0.15 | 0.22±0.10 |
| | 2 | 325.00±16.00 | 13.60±0.95 | 3.15±0.25 | 0.71±0.15 | 0.21± 0.15 |
| | 3 | 327.00±17.00 | 12.72±0.70 | 3.20±0.30 | 0.68±0.13 | 0.24±0.12 |
| | 4 | 315.00±18.00 | 11.50±0.96 | 3.15±0.36 | 0.65±0.10 | 0.17±0.15 |
| | 5 | 337.00±19.00 | 13.60±0.89 | 3.00±0.75 | 0.64±0.16 | 0.19±0.11 |
| **Male** | 1 | 515.00±6.00 | 16.00±0.90 | 4.62±0.86 | 0.85±0.20 | 4.15±0.35 |
| | 2 | 517.00±61.00 | 15.15±0.98 | 4.23±0.79 | 0.90±0.20 | 4.1±0.33 |
| | 3 | 550.00±62.00 | 15.50±0.75 | 4.15±0.56 | 0.87±0.18 | 3.9±0.43 |
| | 4 | 518.00±63.00 | 17.10±0.85 | 4.36±0.81 | 0.82±0.17 | 4.0±0.30 |
| | 5 | 545.00±64.00 | 16.00±0.95 | 4.26±0.75 | 0.97±0.25 | 4.0±0.50 |

Table S11. Effect of DSA-09 on the weight of organs of rats treated for 14 days.

| Sex | Animal No. | Bodyweight before sacrifice | Liver (g/100g) | Kidney (g/100g) | Spleen (g/100g) | Testis /Ovary (g/100g) |
|---|---|---|---|---|---|---|
| **Female** | 1 | 335.00±15.00 | 4.10±0.60 | 0.91±0.11 | 0.19±0.02 | 0.7±0.01 |
| | 2 | 325.00±16.00 | 4.18±0.75 | 0.97±0.09 | 0.22±0.02 | 0.6±0.01 |
| | 3 | 327.00±17.00 | 3.89±0.55 | 0.98±0.13 | 0.21±0.10 | 0.7±0.02 |
| | 4 | 315.00±18.00 | 3.65±0.53 | 1.00±0.80 | 0.21±0.04 | 0.5±0.01 |
| | 5 | 337.00±19.00 | 4.40±0.54 | 0.89±0.11 | 0.19±0.04 | 0.6±0.01 |
| **Male** | 1 | 515.00±60.00 | 3.11±0.47 | 0.9±0.10 | 0.17±0.03 | 0.9±0.06 |
| | 2 | 517.00±61.00 | 2.93±0.44 | 0.82±0.70 | 0.17±0.05 | 0.91±0.08 |
| | 3 | 550.00±62.00 | 2.98±0.55 | 0.82±0.70 | 0.17±0.04 | 0.89±0.04 |
| | 4 | 518.00±63.00 | 3.3±0.43 | 0.84±0.80 | 0.16±0.03 | 0.91±0.09 |
| | 5 | 545.00±64.00 | 2.94±0.48 | 0.78±0.60 | 0.18±0.09 | 0.92±0.07 |

Table S12. Change in the organ to bodyweight ratio (g/100g bodyweight) in the rats treated for 14 days with DSA-09.

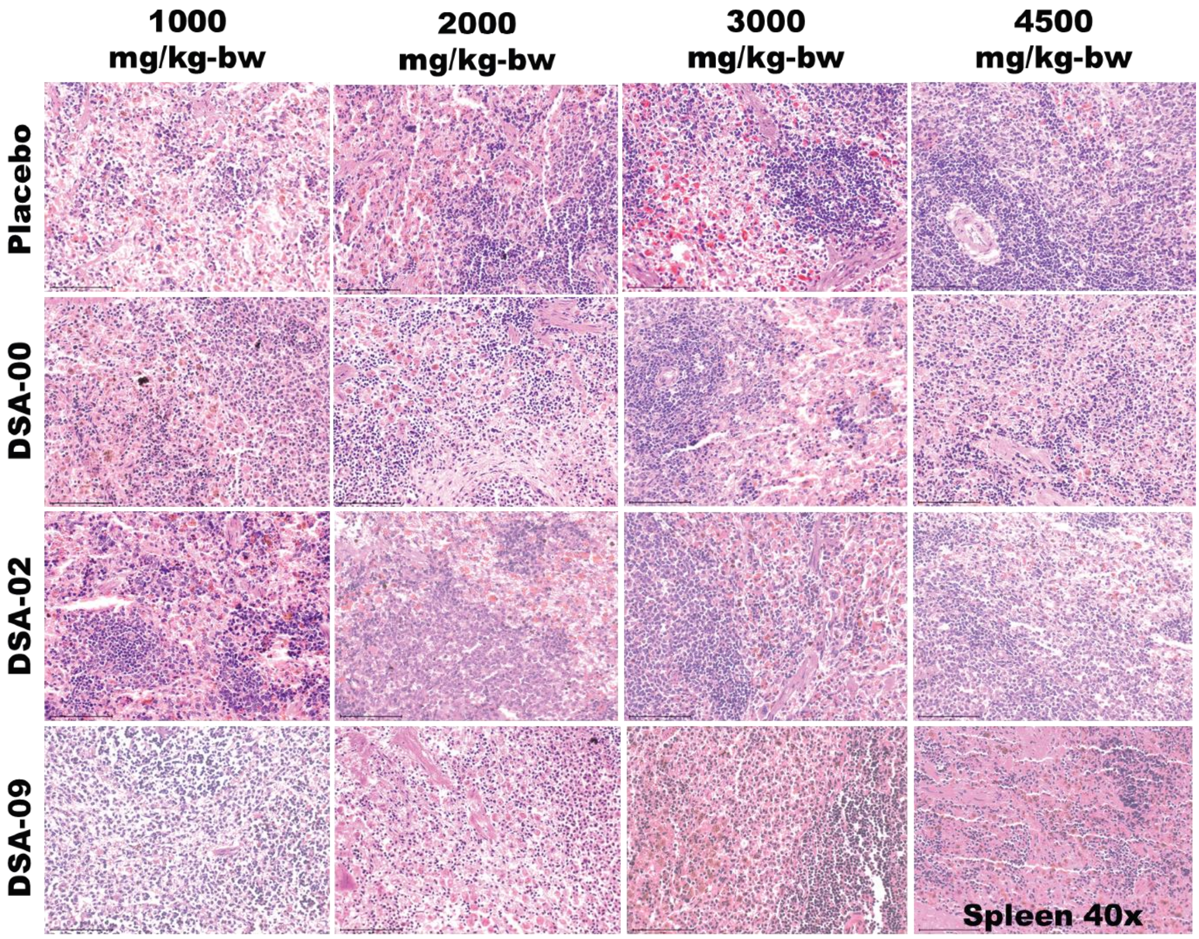


FIGURE S1 The microphotographs (x40) of spleen at 1000, 2000, 3000, 4500 mg/kg-bw groups of thiourea derivatives treated and placebo males and females.

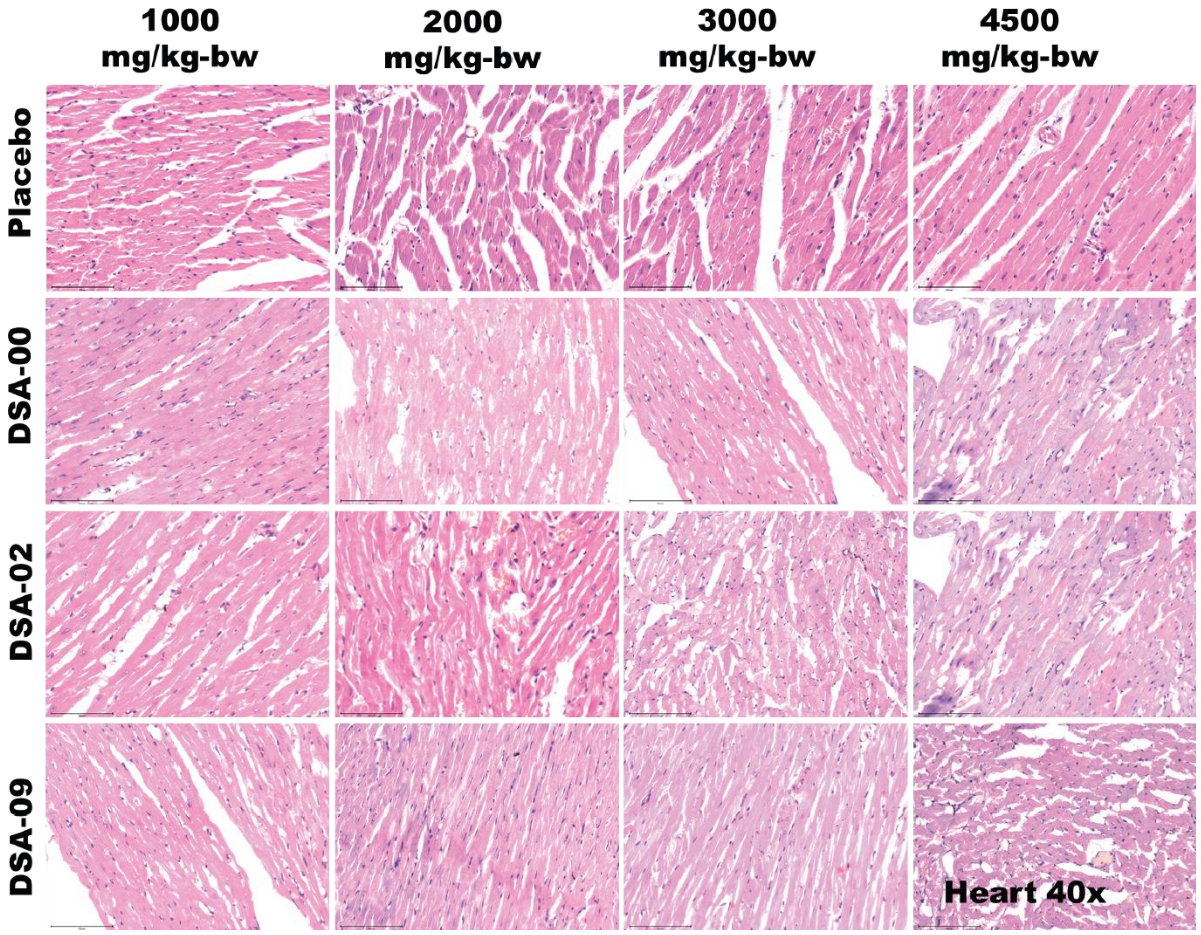


FIGURE S2 The microphotographs (x40) of heart at 1000, 2000, 3000, 4500 mg/kg-bw groups of thiourea derivatives treated and placebo males and females.

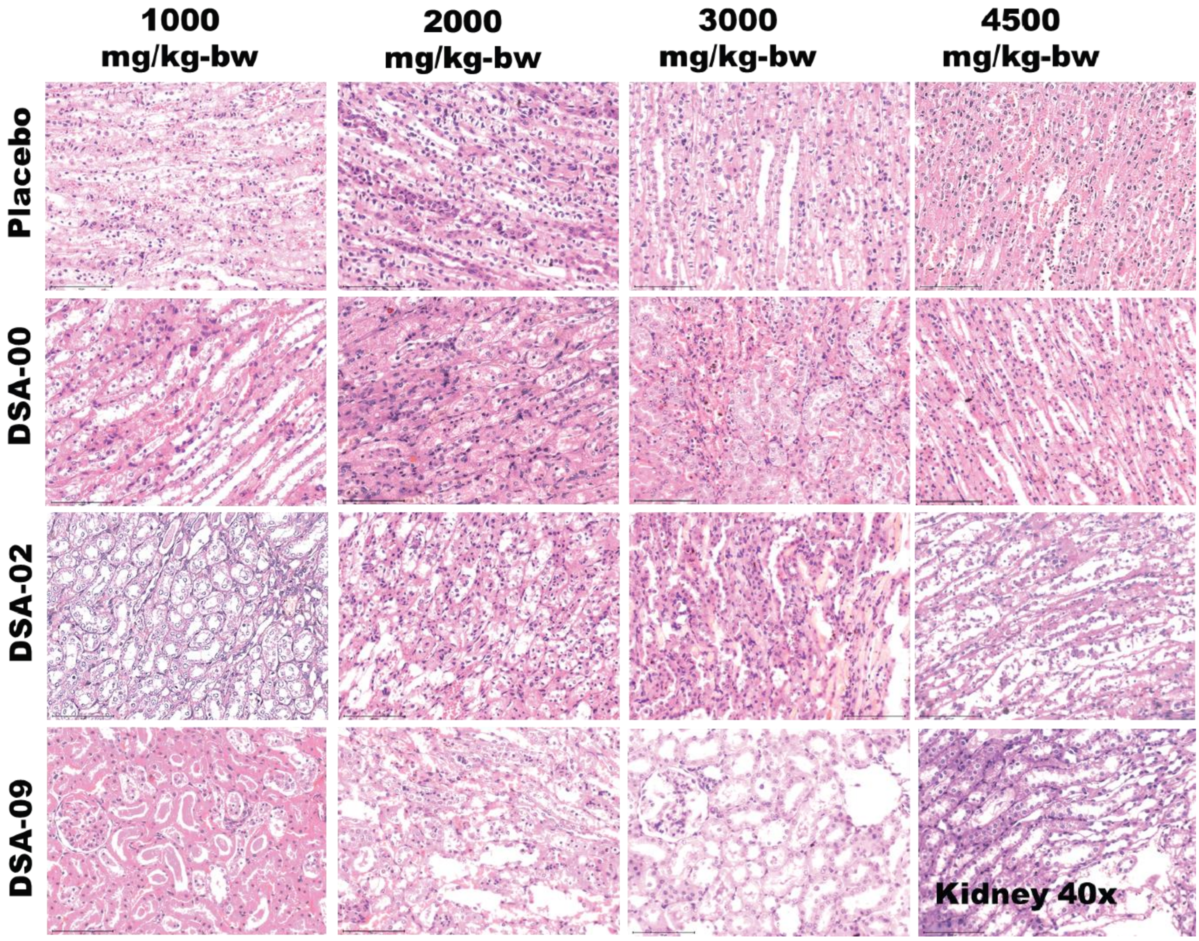

FIGURE S3 The microphotographs (x40) of kidney at 1000, 2000, 3000, 4500 mg/kg-bw groups of thiourea derivatives treated and placebo males and females.

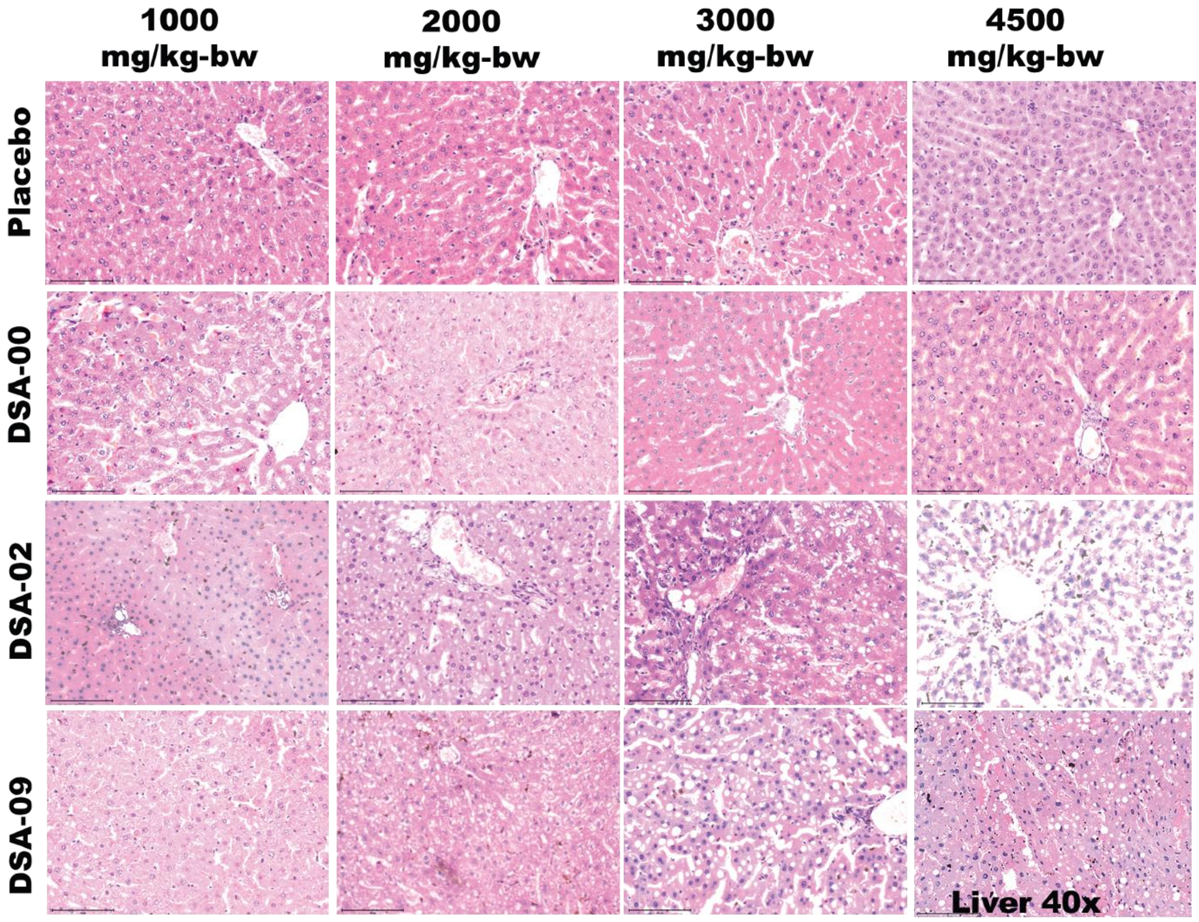


FIGURE S4 The microphotographs (x40) of Liver at 1000, 2000, 3000, 4500 mg/kg-bw groups of thiourea derivatives treated and placebo males and females.

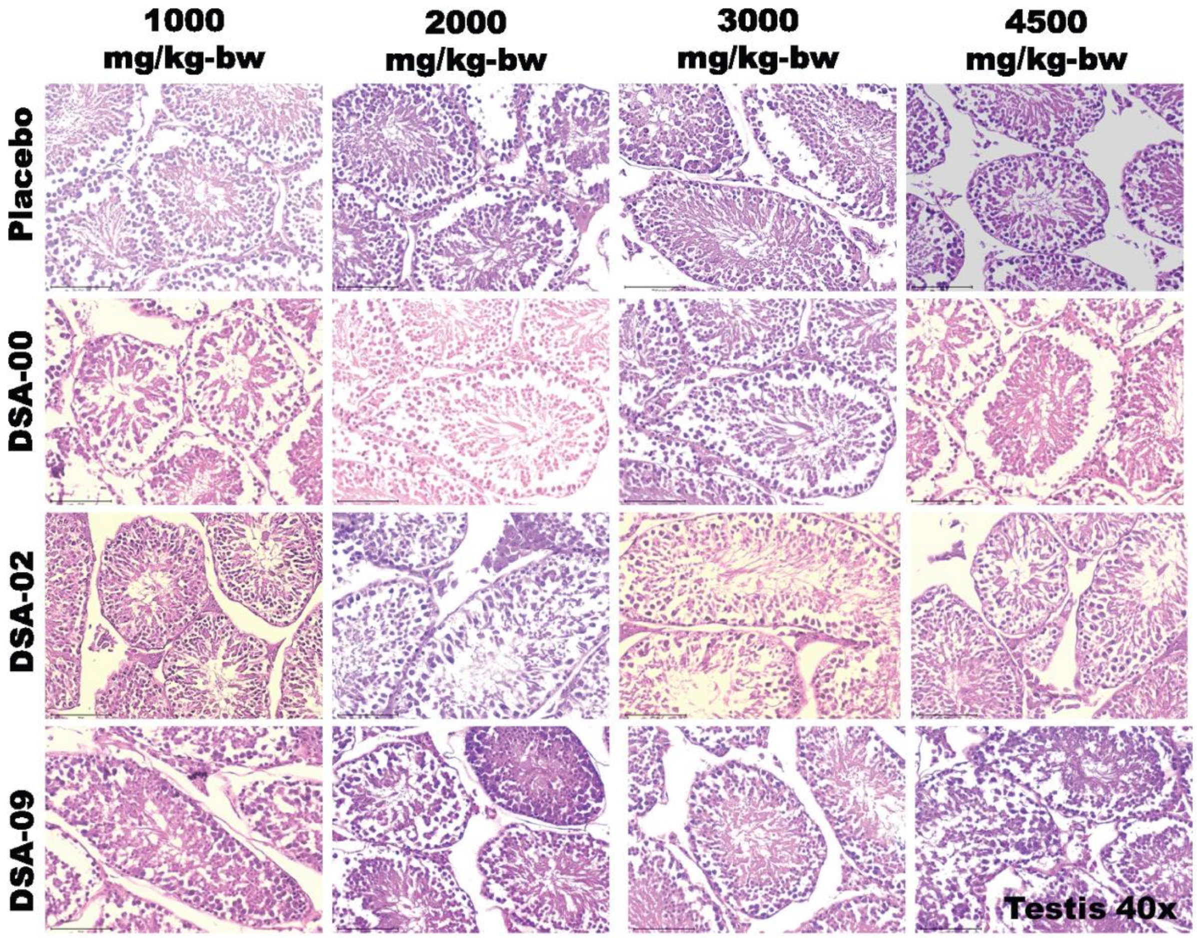


FIGURE S5 The microphotographs (x40) of Testis at 1000, 2000, 3000, 4500 mg/kg-bw groups of thiourea derivatives treated and placebo males.

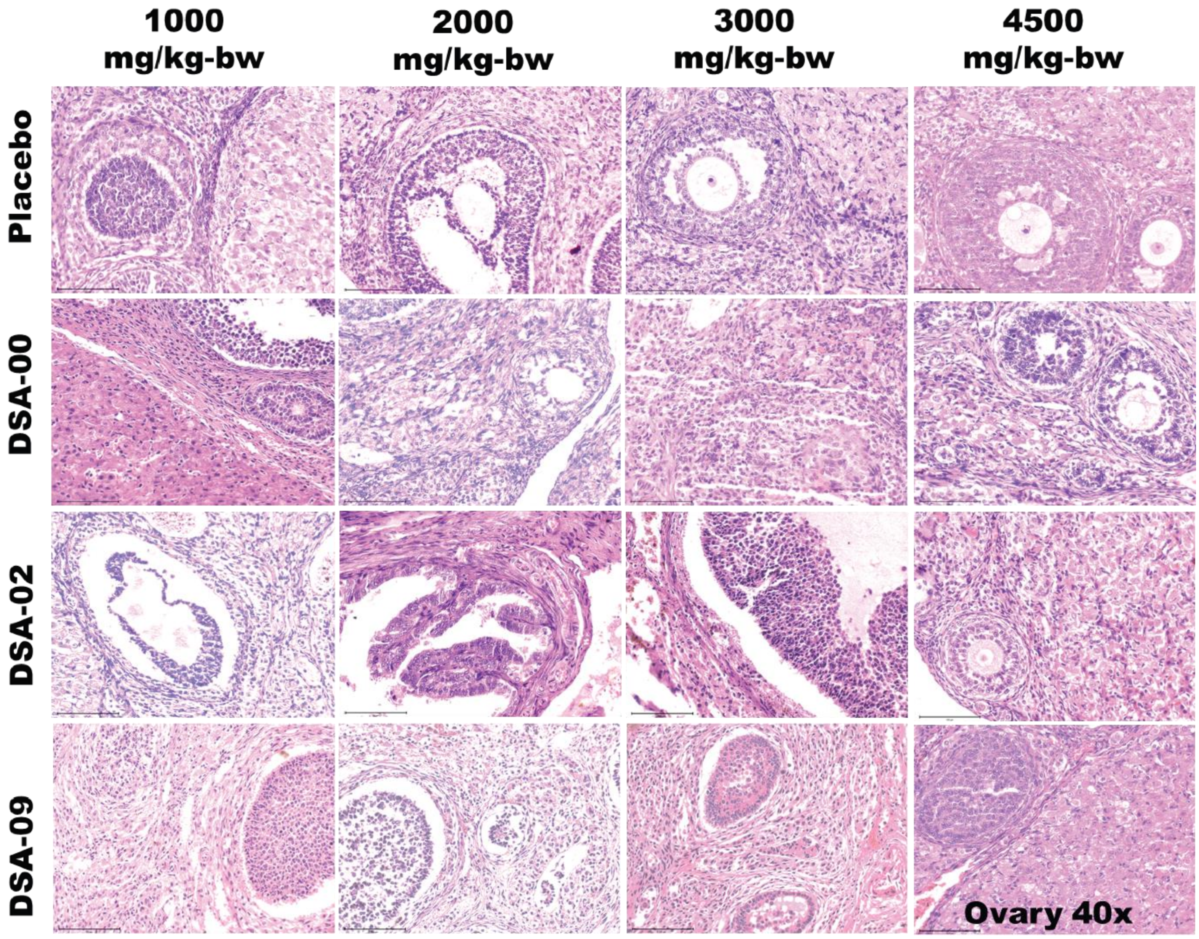


FIGURE S6 The microphotographs (x40) of Ovary at 1000, 2000, 3000, 4500 mg/kg-bw groups of thiourea derivatives treated and placebo females.

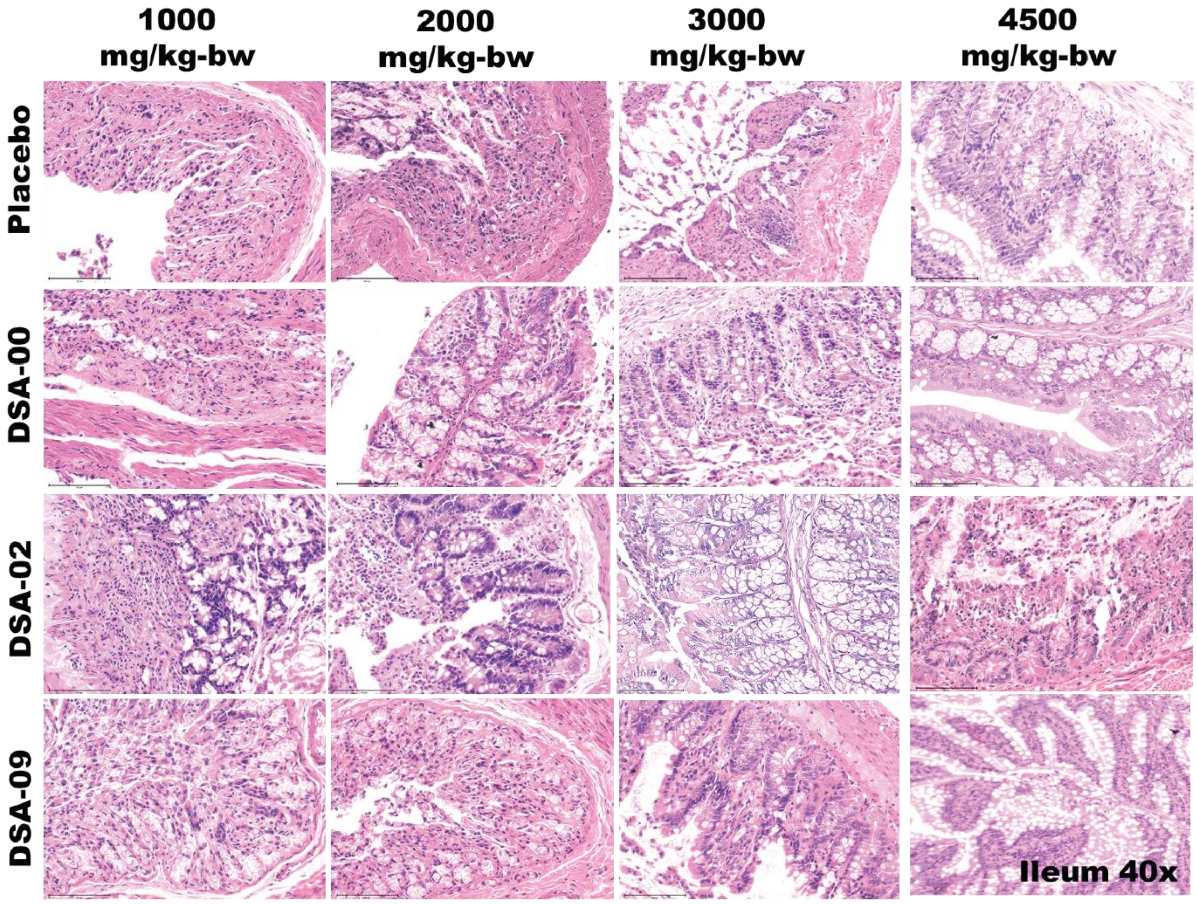


FIGURE S7 The microphotographs (x40) of Ileum at 1000, 2000, 3000, 4500 mg/kg-bw groups of thiourea derivatives treated and placebo males and females.

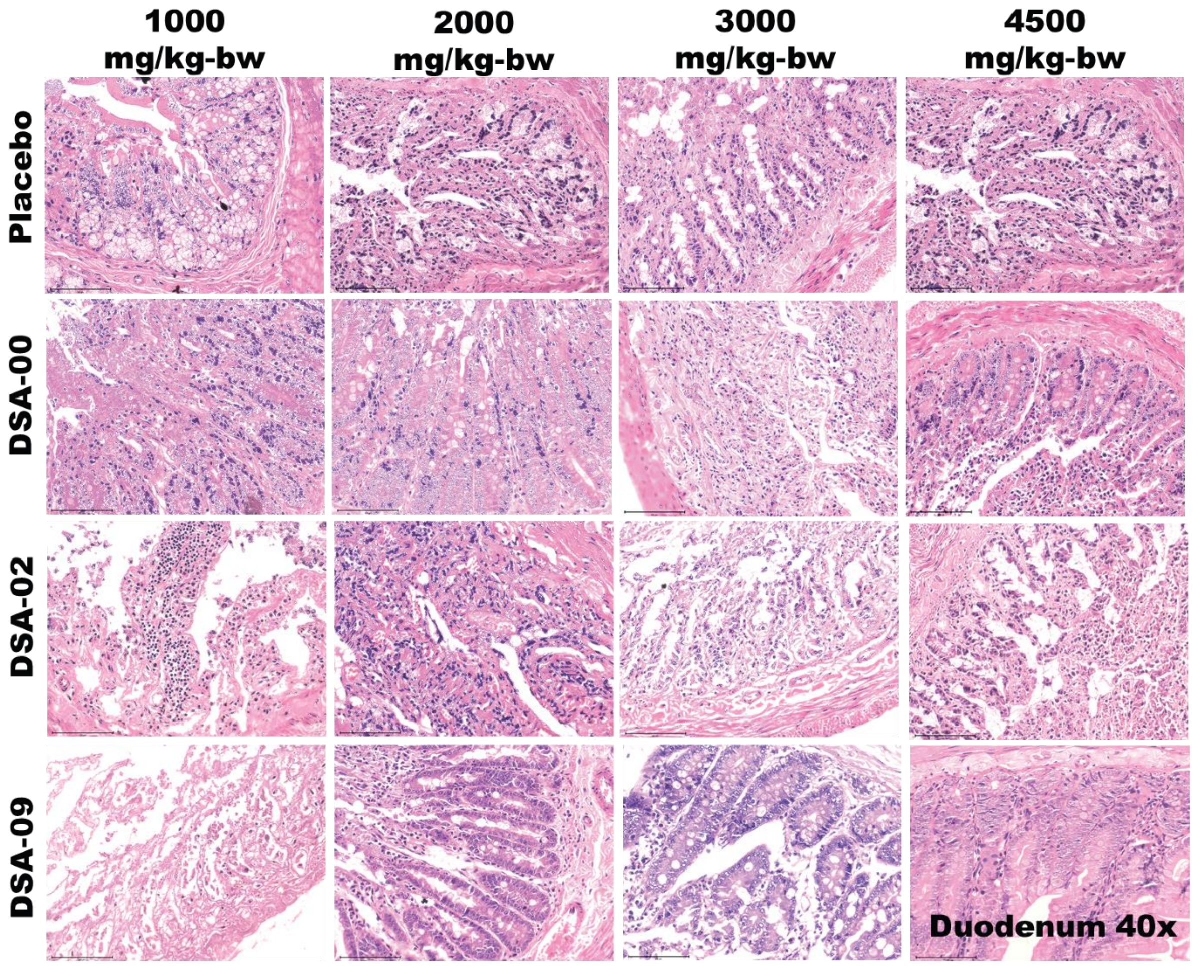


FIGURE S8 The microphotographs (x40) of Duodenum at 1000, 2000, 3000, 4500 mg/kg-bw groups of thiourea derivatives treated and placebo males and females.
